# Structural and Immunological Similarities Between the Metacyclic and Bloodstream Form Variant Surface Glycoproteins of the African Trypanosome

**DOI:** 10.1101/2022.09.28.508705

**Authors:** Monica Chandra, Sara Đaković, Konstantina Foti, Johan Zeelen, Monique van Straaten, Francisco Aresta-Branco, Eliane Tihon, Nicole Lübbehusen, Thomas Ruppert, Lucy Glover, F. Nina Papavasiliou, C. Erec Stebbins

**Affiliations:** Division of Structural Biology of Infection and Immunity, German Cancer Research Center, Heidelberg, Germany; Division of Immune Diversity, German Cancer Research Center, Heidelberg, Germany; Institut Pasteur, Université Paris Cité, Trypanosome Molecular Biology, Department of Parasites and Insect Vectors, F-75015, Paris, France; Centre for Molecular Biology at the University of Heidelberg (ZMBH), DKFZ-ZMBH Alliance, Heidelberg, Germany

## Abstract

During infection of mammalian hosts, African trypanosomes thwart immunity using antigenic variation of the dense Variant Surface Glycoprotein (VSG) coat, accessing a large repertoire of thousands of genes and pseudogenes and switching to antigenically distinct copies. The parasite is transferred to mammalian hosts through the bite of the tsetse fly. In the salivary glands of the fly, the pathogen adopts the metacyclic form and expresses a limited repertoire of VSG genes specific to that developmental stage. It has remained unknown whether the metacyclic VSGs possess distinct properties associated with this particular and discrete phase of the parasite life cycle. We show here using bioinformatic, crystallographic, and immunological analyses of three metacyclic VSGs that they closely mirror the known classes of bloodstream form VSGs both in structure and in the immunological responses they elicit.

## Introduction

*T. brucei* is a unicellular pathogen and the cause of African trypanosomiasis in both human and cattle in sub-Saharan Africa (1,2). Transmitted by the tsetse fly (Glossina sp), *T. brucei* is an extracellular pathogen, which swims freely in the bloodstream and is continuously exposed to immune system surveillance (unlike its relatives, *T. cruzi* and *Leishmania*, which are intracellular pathogens) (3). The bloodstream form (BSF) of *T. brucei* is covered by approximately ten million copies of the **V**ariant **S**urface **G**lycoprotein (**VSG**) molecule, which comprises over 90% of the surface protein of the pathogen (4,5). While there are thousands of VSG-encoding genes and pseudogenes in the trypanosome genome, only one VSG is expressed at a given time from one of numerous bloodstream expression sites (BES) (6). During the course of infection in the mammalian host, the pathogen undergoes antigenic variation, a phenomenon by which cells expressing one particular VSG are cleared by the humoral arm of the host immune system while a sub-population of trypanosomes switch their coat protein to an antigenically distinct variant and thus can no longer be identified and cleared (7). The parasites with the “new” coat escape the host immune system and proliferate. This results in waves of parasitemia that characterize trypanosome infection(8).

The VSG polypeptide can be roughly divided into three functional units: the signal peptide for delivery to the cell surface (and which is cleaved during translocation), a large N-terminal domain (NTD) that forms the bulk of the molecule, and the smaller C-terminal domain (CTD) that is tethered to the NTD by a flexible linker that can often be removed by endogenous and exogenous proteases (5,9). The CTD bears the GPI-anchor, which attaches the VSG to the membrane. The N- and C-terminal domains of the VSGs are divided into different sub-categories based on sequence similarity and the positions of the cysteine residues. Five different classifications of VSG NTDs and six types of the CTD have been suggested amongst other classifications from sequence analysis (10–12). Despite the wide arrays of antigenically distinct VSGs, there are only six atomic-resolution structures of the NTD that have been published: from *T. brucei brucei* VSG1 (MITat 1.1), VSG2 (MITat 1.2 or VSG221), VSG3 (MITat 1.3 or VSG224), VSG13 (MITat 1.13), ILTat1.24, as well as VSGsur from *T. brucei rhodesiense*, with three of these determined in the last four years (13–17). Structures of the smaller, buried, and more flexible CTDs from VSG2 and IlTat1.24 have been determined separately by NMR (18,19).

The published NTD structures show that the tertiary folds of the VSGs share overall conservation, each resembling a dumbbell-shaped entity with upper and lower globular elements (“lobes”) connected by a three-helix bundle core (9). These lobes are found as inserted sequences that connect the helices, replacing short linkers seen in more minimal bundle architectures. The conservation of this overall tertiary fold has suggested that the distinct antigenicity of the VSGs is produced primarily by the divergence in surface amino acids, although the recent structure of VSG3 has broadened the structural “diversity space” of these coat proteins (16). The presence of a post-translational modification (PTM) on VSG3 (an *O*-glycan at the top-surface of VSG NTD) added another layer of complexity to VSG antigenicity, as it is important for host immune evasion. In addition, the more recent structures of VSG13 and VSGsur (17) have shown that many architectural assumptions need reassessment, as these VSGs show significant deviation from the folds of older structures. Finally, the structure of VSG3 and recent biochemical work (20) have suggested that the VSGs of this class likely adopt monomeric (low concentration) and trimeric (high concentration) oligomeric states, in contrast to the strict dimers of other VSGs.

*T. brucei* has a complex life cycle which alternates between the mammalian host and its vector, the tsetse fly. The metacyclic form of *T. brucei* inhabits the salivary gland of the tsetse fly and is injected into the mammalian host when the vector takes a bloodmeal. The metacyclic form is infectious for the mammalian host. Metacyclic cells differentiate further to the rapidly growing bloodstream form soon after delivery into the host. In contrast to bloodstream VSGs, metacyclic VSGs (mVSGs) are expressed from a dedicated monocistronic metacyclic expression site (MES) that is shorter than the expression sites associated genes (ESAGs) containing bloodstream expression sites (BES)(21,22). A specific subset of VSGs is expressed in the metacyclic stage, identified as mVSGs 397, 531, 559, 636, 639, 653, 1954, and 3591 (for the Lister 427 strain, (23,24)). These mVSGs represent the first and primary antigenic surface that is presented to the mammalian host immune system(25–28).

Because of these distinct stages of development and pathogenesis, it has been hypothesized that the mVSGs could manifest differences with bloodstream form VSGs in some manner related to their use in the initial stage of mammalian infection (29). These may include structural divergence as well as differences in post-translational modifications (PTMs), such as are found in VSG3 (16). VSG3 is topologically distinct from all previously characterized VSGs, potentially differing in oligomerization states, and as noted above, harbors *O*-linked hexose chains that are potent immune-modulators. This expansion of the “diversity space” of the VSGs raised the intriguing question: Would the mVSGs show additional structural or chemical diversity that might serve the pathogen in the first stages of entry into the host organism from the tsetse fly?

To begin to address these questions, we have solved the high-resolution crystal structures of mVSG397, mVSG531, and mVSG1954 from *T. brucei brucei*, strain Lister 427. Our results lead us to conclude that, as a whole, the mVSGs closely resemble the bloodstream form surface coat proteins both in structure and function.

## Methods

### Cloning and production of *T. brucei* strains

mVSG397, mVSG531 and mVSG1954 from *T. b. brucei* were cloned into pUC19 the plasmid (BioCat, Germany), and introduced into *T. b. brucei* strain Lister 427 as described below. The mVSGG1954 mutant S321A was generated by site-directed mutagenesis using the QuikChange Lightening kit (Agilent Technologies) according to the manufacturer’s protocol. Transfections were performed into a *T. b. brucei* cell line expressing VSG2, termed 2T1(30). First, 5–10 μg of purified plasmids were mixed with 100 μl of 3 × 10^7^ cells in Tb-BSF transfection buffer (90 mM phosphate buffer, pH 7.3, 5 mM KCl, 50 mM HEPES, pH 7.3, 0.15 mM CaCl_2_) and electroporated using a Lonza Nucleofector, program X-001. After 8 h of incubation, hygromycin B was added to a concentration of 25 μg/ml. Single-cell clones were obtained by serial dilutions in 24-well plates in standard culture medium. The surviving clones were confirmed by sequencing. RNA was isolated using the RNeasy Mini Kit (QIAGEN). The isolated RNA was treated with DNase in DNase Turbo Kit (Invitrogen) according to the manufacturer’s protocol. Complementary DNA was synthesized with ProtoScript Reverse Transcriptase (NEB) according to the manufacturer’s protocol. Amplification was performed with a forward primer binding the spliced leader sequence and a reverse primer binding in the VSG 3’-untranslated region (UTR) using Phusion high-fidelity DNA polymerase. PCR products were purified by gel extraction from a 1% agarose gel, followed by the NucleoSpin Gel and PCR clean-up kit (Macherey-Nagel), and sent to Eurofins (Ebersberg) for sequencing using the same primers as for the PCR.

### Purification and Crystallization of mVSG397, mVSG531, and mVSG1954

*T. b. brucei* strains expressing mVSG397, mVSG531 and mVSG154 were cultivated *in vitro* in HMI-9 medium (formulated as described (17)), supplemented with 10% fetal calf serum (Gibco), L-cysteine and β-mercaptoethanol. Cells were cultured at 37 °C and 5% CO_2_. VSGs were purified according to established protocols (31). Briefly, cells were pelleted and then lysed in 0.2 mM ZnCl_2_. The lysis mixture was centrifuged and the pellet containing the membrane material was resuspended in prewarmed (42 °C) 20 mM HEPES pH 7.5, 150 mM NaCl. Following a second centrifugation, supernatant containing VSG protein was loaded onto an anion-exchange column (Q-Sepharose Fast-Flow, GE Healthcare), which had been equilibrated with 20 mM HEPES pH 7.5, 150 mM NaCl. The flow-through and washes fractions containing highly pure VSG was concentrated in an Amicon Stirred Cell with 10 kDa MWCO membrane. The concentrated VSGs were subjected to size exclusion chromatography as the last clean-up step using HiLoad 16/600 Superdex 200 column (GE Healthcare) in AKTA pure chromatography system (GE Healthcare).

Concentrated mVSG397 was digested with trypsin in a 1:100 trypsin:VSG ratio on ice for 1 hour. The mVSG397 NTD was purified using a gel filtration chromatography Superdex 200 Increase 10/300GL column (GE Healthcare) equilibrated with 10 mM Hepes-NaOH, 150 mM NaCl. The fractions containing the protein were concentrated to 5 mg/ml. Crystals were grown by vapour diffusion using 100 mM SPG buffer (pH 5.2) and 21 % (W/V) PEG 1500 and soaked in 100 mM SPG buffer (pH 5.2), 21 % (W/V) PEG 1500 and 10 % (W/V) PEG 400 before flash cooling to 100 K (−173.15 °C). For phasing, crystals were grown by vapour diffusion using 100 mM SPG buffer (pH 5.2) 24 % (W/V) PEG 2000 MME and shortly soaked in 100 mM SPG buffer (pH 5.2), 24 % (W/V) PEG 200 MME, 50 mM KI and 25 % (W/V) ethylene glycol before flash cooling to 100 K (−173.15 °C). Data sets were collected at the Paul Scherrer Institut, Swiss Light Source, Villingen, at a wavelength of 1.0 Å or 2.066 Å (native and KI soaked crystals, respectively). The structure was solved using single wavelength anomalous diffraction (SAD) with the software CCP4I2-CRANK2 (32), and an initial model was built with Buccaneer(33). This model was used for molecular replacement in PHENIX (34,35) against the higher resolution native data. The structure was refined and built using PHENIX, COOT (36) and PDB-REDO (37).

Native crystals of mVSG531 were grown by vapor diffusion using hanging drops formed from mixing a 1:1 volume ratio of the full-length protein with an equilibration buffer consisting of 25% PEG 1500, 0.1 MMT buffer pH 6.5, 3% glucose. Crystals of only the NTD appeared after 3 weeks at room temperature. For cryoprotection, crystals were transferred directly into a buffer with 10% glycerol and flash-cooled immediately afterward to 100 K (−173.15 °C). Crystals formed in the space group P2_1_ with four dimers of VSG531 in the asymmetric unit (total of eight independent copies of the VSG531 polypeptide). Data were collected at the European Synchrotron Radiation Facility (ESRF) in Grenoble at beamline ID29 and processed onsite through the EDNA framework Fast Processing System (PMID: 19844027). The structure was solved by molecular replacement using the model of VSG1 (MITat1.1, PDB ID 5LY9) with PHASER as implemented in the PHENIX package(38). Iterative cycles of model building with the PHENIX (39), manual adjustment in COOT, and refinement in PHENIX let to a final model with and R/Rfree of 21.34%/24.38% with no Ramachandran outliers. An N-linked carbohydrate is present in each monomer at residue N295.

The concentrated mVSG1954 was subjected to limited proteolysis using trypsin in 1:50 trypsin to VSG ratio at 4 ºC for 3 hours to remove the more flexible C-terminal domain. The proteolysis reaction was stopped by adding phenylmethylsulfonylfluoride (PMSF) to 1 mM final concentration. The N- and C-terminal domains were separated by size exclusion chromatography using HiLoad 16/600 Superdex 200 column (GE Healthcare) in AKTA pure chromatography system (GE Healthcare). Native crystals were grown by vapor diffusion using sitting drops formed from mixing a 1:1 volume ratio of the protein with an equilibration buffer consisting of 20% PEG 3350, 0.4M NaCl at the University of Heidelberg Crystallization Platform. Crystals appeared after 19 hours at 18 °C. The native crystals diffracted to 2.28 Å at Paul Scherrer Institut, Swiss Light Source, Villigen, Switzerland. The derivative crystals were grown by vapor diffusion using sitting drops formed from mixing a 1:1 volume ratio of the protein with an equilibration buffer consisting of 25% PEG 1500, 0.4M NaBr for halide phasing. The crystals appeared after 3 days following incubation at 22 °C. For cryoprotection, crystals were transferred directly into a stabilization buffer with 0.75M NaBr and 20% glycerol and flash-cooled immediately afterward to 100 K (−173.15 °C). Anomalous X-ray diffraction data were collected at 0.9198 Å wavelength to obtain single-wavelength anomalous dispersion (SAD) dataset. The best diffracting derivative crystal diffracted to 1.68 Å. Crystals formed in the space group P321 with a monomer of mVSG1954 in the asymmetric unit. The substructure atoms were found using SHELX(40) in the CCP4i2 (41) crystallography software suite. Initial model building was performed in Arp/wArp (42) and BUCCANEER in the same suite. Subsequently, the model was further refined in PHENIX. However, after data quality analysis, the last 300 frames in the dataset were opted out from the processing in xia2/DIALS(43–45) to avoid incorporating data with radiation damage. The refined structure was used to perform molecular replacement with PHASER in PHENIX on the new reprocessed data. Subsequent refinement was performed in PHENIX. Cycles of manual model examination using COOT and improvement with automated refinement completed the model. Final model with and R/Rfree of 20.9%/21.3% with no Ramachandran outliers. An N-linked carbohydrate is present in each monomer at residue N374.

### Suramin Resistance Assays

Performed as previously described (17).

### Mass spectrometry with purified mVSG1954 wt and S321A

Protein samples were loaded on a self-packed reversed-phase column (0.8 × 2 mm, Poros R1) for desalting and concentration using 0.3% formic acid (0.3 ml/min). After 3 minutes proteins were eluted with 40% isopropanol, 5% acetonitrile, 0.3% formic acid (0.04ml/min) and analyzed with a QTof mass spectrometer (maXis, Bruker Daltonics) after electrospray ionization. Data were deconvoluted using the ESI Compass 1.3 Maximum Entropy Deconvolution Option (Bruker) to determine the molecular weight of the proteins.

### Growth of Metacyclic Cells

*T. brucei* PCF 29-13 cells were transfected by plasmid that encodes for tetracycline-inducible RBP6 overexpression (kindly gifted by Lucy Glover, Institute Pasteur, Paris, France, (Tihon et al., 2022)). The cells were cultured in SDM-80 media (SDM-79 without glucose). SDM-80 media was supplemented with 7.5 μg/ml Hemin, 10% heat-inactivated FBS, and 50 mM N-Acetylglucosamine. Procyclic cells were grown at 27°C with 5% CO_2_. On day 0, RBP6 overexpression was induced with 10 μg/μl tetracycline at 2 × 10^6^ cell density. The growth of both induced and uninduced cultures were examined and diluted to 2 × 10^6^ cells per ml. Tetracycline was added every day during differentiation. The occurrence of MCF was examined using flow cytometry on day 6 by detecting the loss of GPEET and EP expression.

### Generation of *α*-mVSG531 and *α*-mVSG1954 polyclonal antisera

The infection procedure was adapted from Aresta-Branco et al., 2019. Antisera against BSF expressing mVSG531 or mVSG1954 (α-mVSG531 and α-mVSG1954) was generated by infecting two C57BL/6J female mice. On day 0, two C57BL/6J female mice were injected (i.p.) by 1000 parasites (BSF expressing either mVSG531 or mVSG1954) suspended in 200 μl HMI-9 media. Berenil (1.25 μg/ml) was administered to both mice on day 4 and 5 post infection (i.p.).

Both mice were sacrificed on day 8 by CO_2_ asphyxiation. Blood was collected from both mice by cardiac puncture. The antisera were separated from the other cellular component by centrifugation in BD Microtainer blood collection tubes. The specificity of the antisera was further confirmed by flow cytometry to stain the original cell line used for the infection.

### Magnetic separation of MCFs and epimastigotes from undifferentiated PCFs

Undifferentiated PCF cells were separated from the non-PCF cells by negative selection using magnetic beads. On day 6 post-induction of RBP6 overexpression. 1 × 10^8^ cells in induced culture were collected by centrifugation at 1800 xg for 10 minutes at 4°C (TX-750 rotor, Thermo Scientific). Monoclonal α-GPEET and α-EP IgG (Cedarlane Labs) were used in 1:50 antibody to cells volume as the primary staining. MACS Goat α-mouse IgG magnetic beads (Miltenyi) was used as the second staining step. To separate the undifferentiated PCF cells, the cell suspension was pass through LS-column placed in MACS magnetic field (Miltenyi), and flow through that contain the non-PCF cells was collected. The column was subsequently washed with 3 × 3 ml of SDM-80 media to collect all the non-PCF cells.

### FACS Analysis

Flow cytometry was performed to detect MCF cells that express mVSG531 or mVSG1954 within the differentiated cells. The cell density from both uninduced and tetracycline-induced culture were counted. Two million cells were used in each flow cytometry sample. The cells were harvested by centrifugation at 5,400 xg for 4 minutes at 4 ºC. Monoclonal α-GPEET and α-EP IgG were conjugated with Allophycocyanin (APC) fluorescence molecule by APC Conjugation Kit – Lightning Link (Abcam). Cells were stained with α-GPEET or α-EP or both α-GPEET and α-EP to detect the loss of GPEET and EP as the indicator of the presence of non-PCF cells in the induced culture. The cells were stained in 1:100 antibody to medium ratio (α-GPEET-APC, α-EP-APC, or both α-GPEET-APC and α-EP-APC) prior to analysis using BD-FACS Calibur Flow Cytometry. 50,000 cells were analyzed from each sample.

Two million undifferentiated PCF cells were mixed with 50,000 BSF cells expressing either mVSG531 or mVSG1954. These samples were used as a positive control that α-mVSG531 and α-mVSG1954 are able to recognize a small population of cells expressing mVSG531 or mVSG1954 in a mixture with other cells. Two million purified differentiated cells were also used in each flow cytometry sample. The differentiated cells were stained by α-mVSG531, α-mVSG1954, or both α-mVSG531 and α-mVSG1954. The cells were harvested by centrifugation at 5,400 x g for 4 minutes at 4 ºC. The cells were stained with the respective antibody in 1:100 antibody to medium ratio (α-mVSG531, α-mVSG1954, or both α-mVSG531 and α-mVSG1954). The cells were subsequently counterstained with 1:1000 α-Mouse IgM-FITC. The cells were analyzed by BD-FACS Calibur Flow Cytometry. 50,000 cells were analyzed from each sample.

## Results

### Structure-focused Classification of Metacyclic and Bloodstream VSGs

Using structural topology as a guide, the known VSG NTD structures can be divided into two “super-families” that correlate well with previous divisions based on primary sequence analyses (Classes A and B), while also providing an architectural explanation for the class division. Class A includes the largest set of members that have been structurally characterized: VSG1, VSG2, ILTat1.24, VSG13, and VSGsur (Fig 1, A and B). This class possesses bottom lobes of globular secondary structure (near the membrane proximal portion of the 3-helix bundle of the VSG NTD; Fig 1A) that are composed of primary sequence that is located close to the C-terminal portion of the NTD. In contrast, Class B is topologically distinct, possessing a bottom lobe that is comprised of sequence from the N-terminal portion of the molecule (Fig 1). This suggests that while there can be many subdivisions within each of these clans, there is a broad overarching bifurcation of the VSGs by protein structural topology that is dictated by the position of the amino acid sequence encoding the bottom lobe structure.

**Fig. 1:**
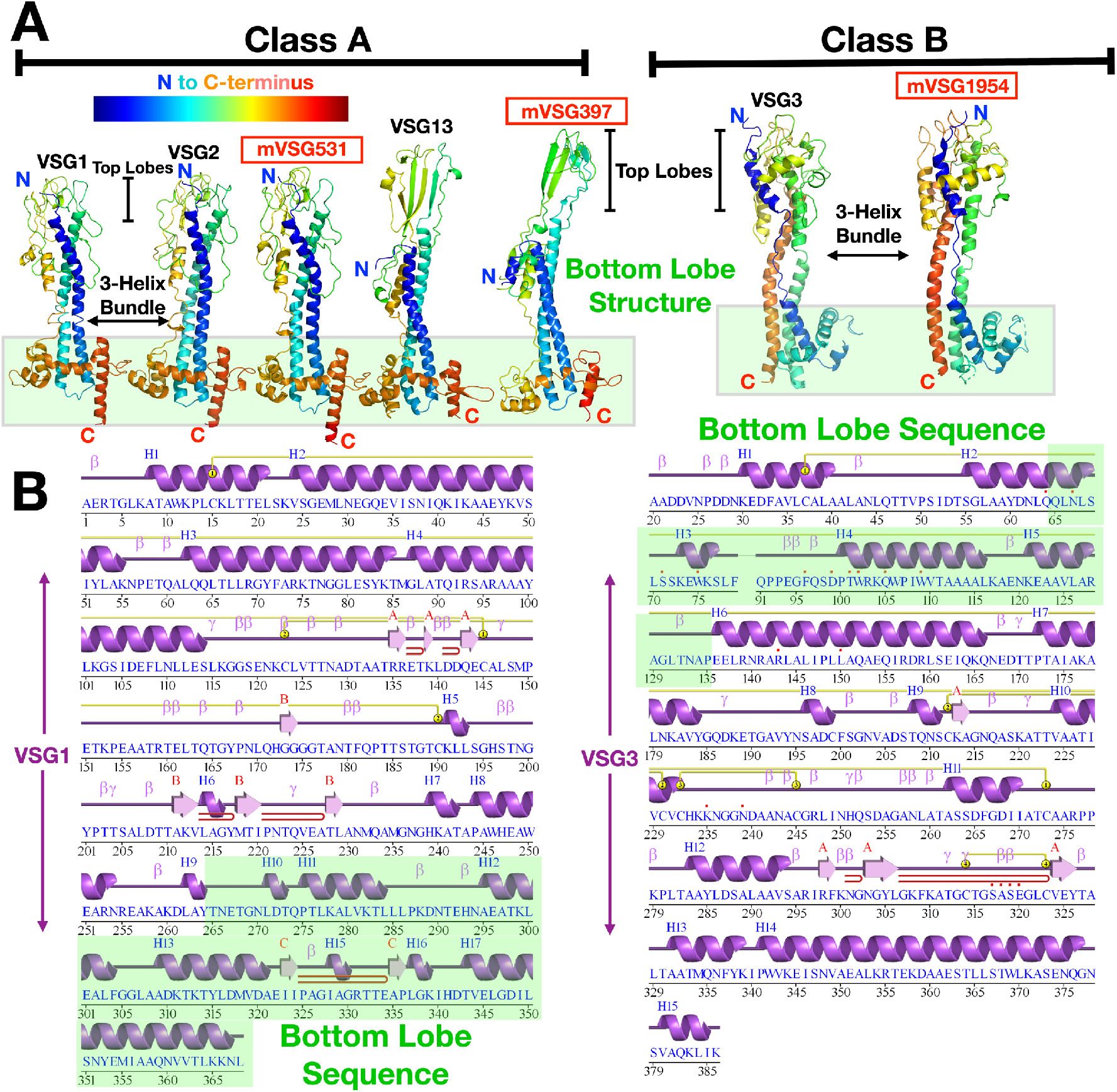
Structural classification of VSG proteins. (A) Two broad superfamily classes of VSGs identified through sequence analysis and defined by structural topology are shown here with representative structures. VSG monomers are shown as ribbon diagrams colored in a gradient from blue to red from N- to C-terminus. Structures reported in this manuscript are denoted in red and described in detail below. The lower, bottom lobe subdomains of the VSGs are highlighted in a green-tinted box. Structures drawn with CCP4*mg* (58). (B) Sequence and structurally-determined secondary structure of two representative VSGs from Class A (VSG1) and Class B (VSG2). The sequence region that forms the bottom lobe is indicated by green highlighting. Secondary structure illustrated with PDBSUM (59).

As a subclass, the metacyclic VSGs studied here do not deviate from this pattern. VSGs 397 and 531 fall within Class A by homology analysis, whereas VSG1954 is a member of Class B. More specifically, by using the classification system outlined in Cross et al. (10), mVSG397 belongs to Class A1 (similar to VSG522, which shares 72% sequence identity with *Trypanosoma brucei rhodesiense* VSGsur (GenBank: ATI14856), whose unusual structure was recently determined (17)), mVSG531 to Class A2 (the same as VSG1 and VSG2), and VSG1954 to Class B.

### Crystal Structure of mVSG397 Shows Close Homology to *T*.*b rhodesiense* BSF VSGsur

To extend the comparison between the mVSGs and the BSF VSGs beyond sequence analysis, we determined the crystal structure of Class A1 mVSG397 to 1.26Å resolution (Methods, Fig 2, Supplementary Fig 1A, Supplementary Table S1). As noted, sequence clustering analysis classified this VSG similarly to VSGsur, and the structure confirms the architectural similarity. Like VSGsur, mVSG397 possesses several distinct conformational features strongly distinguishing it from known structures of both Class A2 and Class B VSGs, while retaining the bottom lobe topological relationship with those Class A structures. A prominent distinction is the top lobe of the VSG, which consists of a large, globular subdomain - a tightly twisted β-sheet in the monomer that forms a β-sandwich in the dimer (Fig 2, A and B). Secondly, unlike the VSG1, VSG2, and VSG3 which possess N-linked glycans at the bottom lobes, the sugar in mVSG397 is positioned similarly to VSGsur, located directly below the top lobe, nearly two-thirds the distance from the bottom of the NTD. A third similarity to VSGsur is the disulfide bond distribution throughout the length of the NTD, whereas the other known VSGs contain such bonds clustered in the upper portion of the top lobe. Finally, as with VSGsur, the N-terminus of the protein is not at the surface, but located below the top-lobe and N-linked glycan, possibly buried significantly when packed on the membrane. Overall, mVSG397 and VSGsur align with a root mean square deviation (RMSD) of 2.33 Å over 334 residues (Fig 2B). Consistent with the functional VSGs possessing antigenic distinctiveness to the immune system, the molecular surface of mVSG397 differs appreciably from that of VSGsur in topography, electrostatic and hydrophobic properties (Fig 2C).

**Fig 2:**
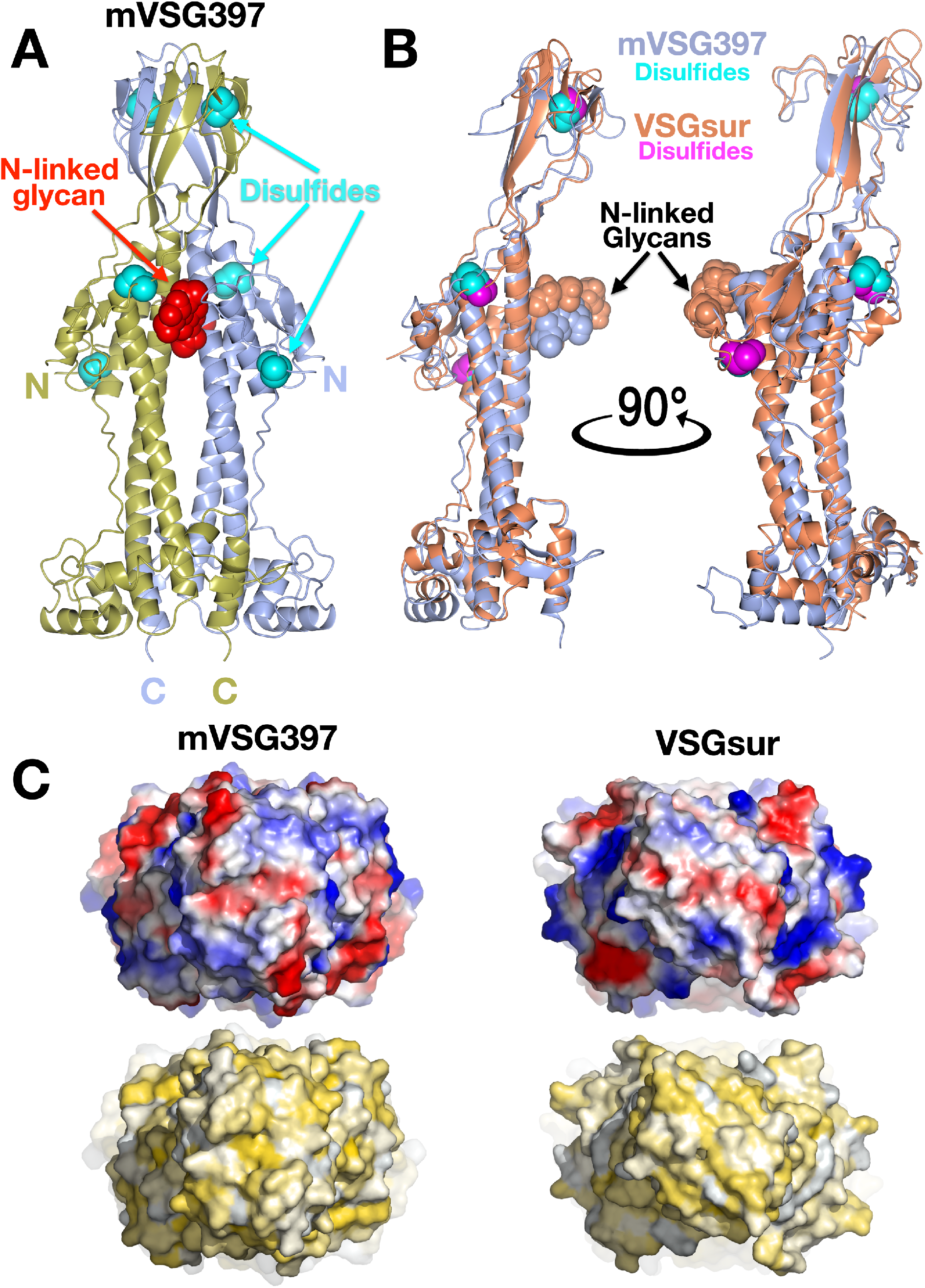
Crystal structure of mVSG397. (A) Ribbon diagram of the (crystallographic) dimer of mVSG397, the individual monomers colored gold and light blue. The N-linked glycans are displayed as red space-filling atoms and disulfide bonds are shown in cyan. ‘N’ and ‘C’ indicate the N and C termini of each monomer. Structures drawn with CCP4*mg* (58) (B) Structural alignment of monomers of VSGsur (orange) and mVSG397 (light blue) with corresponding glycans and disulfides shown in space filling representation, the sugars colored the same as the protein to which they are linked, the disulfides in cyan (mVSG397) and magenta (VSGsur). Structures drawn with CCP4*mg* (58) (C) 90-degree rotation of the structures in (A) and (B) to view the “top” of the VSGs, rendered as molecular surfaces. Top row shows the surface colored by electrostatic potential (red indicating acidic/negatively charged, blue indicating basic/positively charged, and white neutral). The bottom row is colored by the Eisenberg hydrophobicity scale (Eisenberg, D., Schwarz, E., Komaromy, M. & Wall, R. Analysis of membrane and surface protein sequences with the hydrophobic moment plot. J. Mol. Biol. 179, 125–142 (1984)), where yellow indicates hydrophobic and white polar. Molecular surfaces are illustrated with PyMOL (60).

The structure of VSGsur was published together with the structure of VSG13, the latter a VSG also possessing a large β-sandwich top lobe, an N-linked glycan directly underneath this subdomain, and disulfides distributed over the length of the NTD. However, the β-sheet domains of VSGsur and VSG13 do not align well (that of VSG13 being less twisted and much broader), and mVSG397 resembles VSGsur in this region and not VSG13. This is consistent with the sequence clustering classification of these A-type VSGs into different subgroups, A1 for both VSGsur and mVSG397 but A3 for VSG13 (10).

Finally, it should be noted that variant VSGsur from *T. b. rhodesiense* binds to the trypanolytic drug suramin, thereby conferring heighted resistance to the drug (17). While mVSG397 possesses less than 30% sequence identity with VSGsur in the NTD, the question arises of whether it could bind suramin as well. Suramin binds VSGsur in a large cavity formed between the helices of the three-helix bundle, directly below the N-linked glycan. However, no such large cavity exists in mVSG397 (Supplementary Fig 2A). Indeed, trypanosome strains expressing VSG397 are not resistant to suramin like VSGsur (Supplementary Fig 2B). A better candidate for suramin binding in the Lister 427 strain would be VSG522, which possesses 72% sequence identity with VSGsur, including key binding residues to the drug, and other studies could examine this possibility.

### Crystal Structure of VSG531 Shows Close Homology to Class A2 Members

We also crystallized mVSG531, predicted to be of the Class A2 group, determining its structure to 1.95Å resolution (Fig 3, Supplementary Figs 1B, Supplementary Table S1, and Methods). Consistent with bioinformatic analyses, mVSG531 is very similar to the structures of other Class A2 members solved from BSF VSGs, particularly VSG1 and VSG2, aligning to each with a RMSD of 2.1Å over 354 residues and 2.4Å over 347 residues, respectively (Fig 3A, (47,48)). In fact, with 35% sequence identity, the structure of VSG1 is similar enough to mVSG531 that we were able to use it to phase the diffraction data through molecular replacement (see Methods, Supplementary Fig 3). Most of the divergence between VSG531 and VSG1/VSG2 occurs in the top lobes, with the bottom lobe showing higher conservation, and the three-helix bundle the most conserved region of the molecule (RMSD close to 1Å, Supplementary Fig 4).

**Fig 3:**
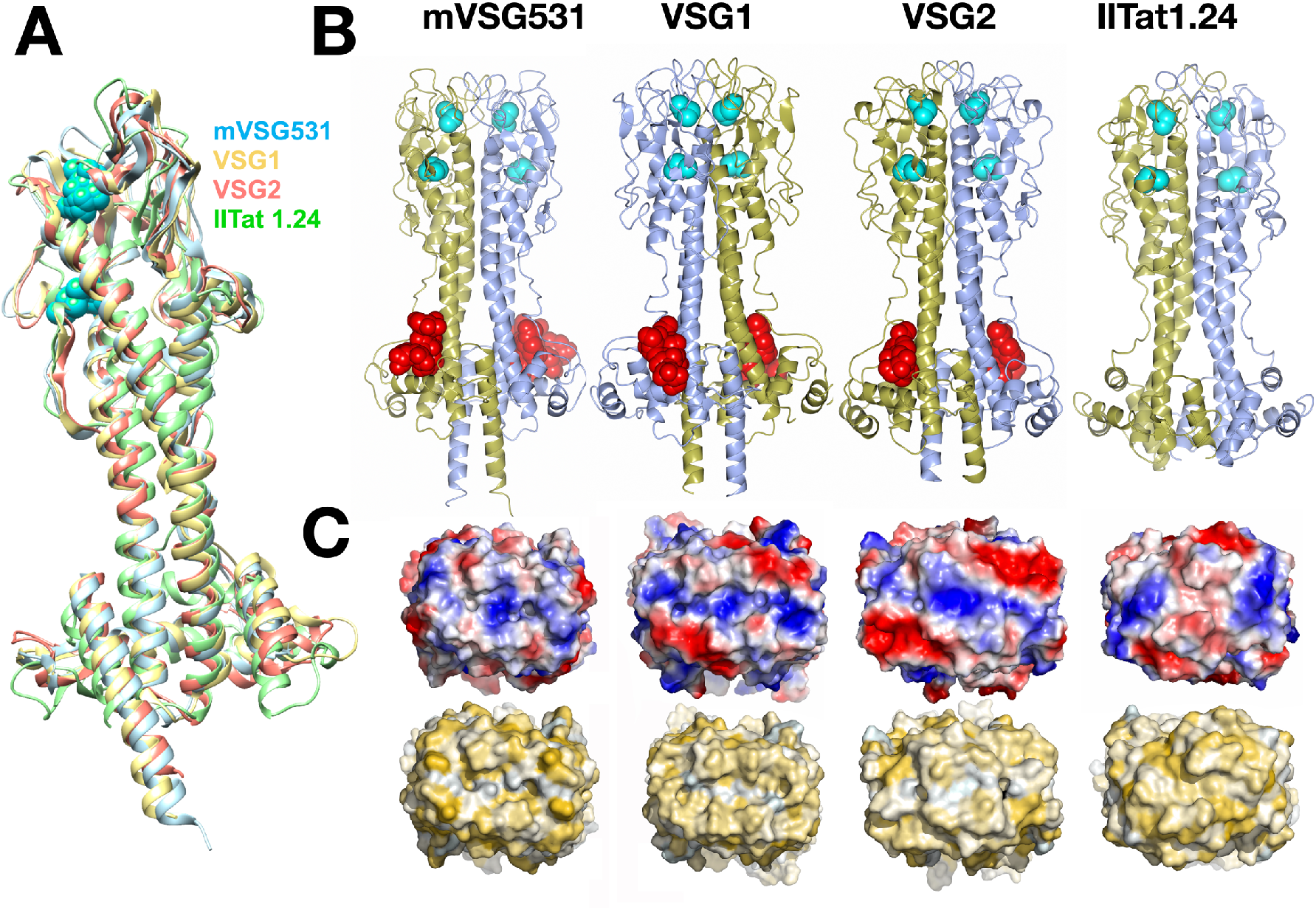
Crystal structure of mVSG531. (A) Superposition of class A2/N2 monomers of mVSG531, VSG1, VSG2, and IlTat1.24 produced using DeepAlign in the RaptorX structure alignment server (47,48). Images of protein structures were generated and edited using CHIMERA (61). (B) Side-by-side comparison of different class A VSG dimer highlighting common fold, disulfide placement (cyan spheres), and N-linked glycan (red spheres). Structures drawn with CCP4*mg* (58). (C) 90-degree rotation of the structures in (B) to view the “top” surface of the VSG, rendered as a molecular surface. Top row shows the surface colored by electrostatic potential (red indicating acidic/negatively charged, blue indicating basic/positively charged, and white neutral). The bottom row is colored by the Eisenberg hydrophobicity scale(62), where yellow indicates hydrophobic and white polar. Molecular surfaces are illustrated with PyMOL (60).

Similarities to VSG1 and VSG2 extend beyond the overall conservation in the fold. The positions of the two cysteine disulfides bonds in the top lobe of the N-terminal domain closely overlap with several of those in structures of bloodstream Class A2 members (Fig 3, A and B), although this does not hold for all Class A members, such as A3 member VSG13 and, as discussed above, Class A1 mVSG397 and VSGsur. The N-linked sugars of VSG1, VSG2, and mVSG531 are located in very similar positions in the lower lobe as well (Fig 3B). Finally, the molecular surfaces of different VSGs show high variance in shape, charge distribution, and polar/hydrophobic character (Figs 2C and 3C), regardless of which subgroup to which they belong.

### Crystal Structure of mVSG1954 Shows Close Homology to class B members

To further compare mVSGs and BSF VSGs, we characterized the structure of Class B member mVSG1954 at 1.68 Å resolution (Methods, Supplementary Fig 1C, Fig 4). mVSG1954 and VSG3 (also class B) show a good degree of structural conservation, aligning with an RMSD of 4.6Å over 308 residues). The conservation extends to the location of the four cysteine disulfides bridges concentrated at the top-lobe of the monomer.

**Fig 4:**
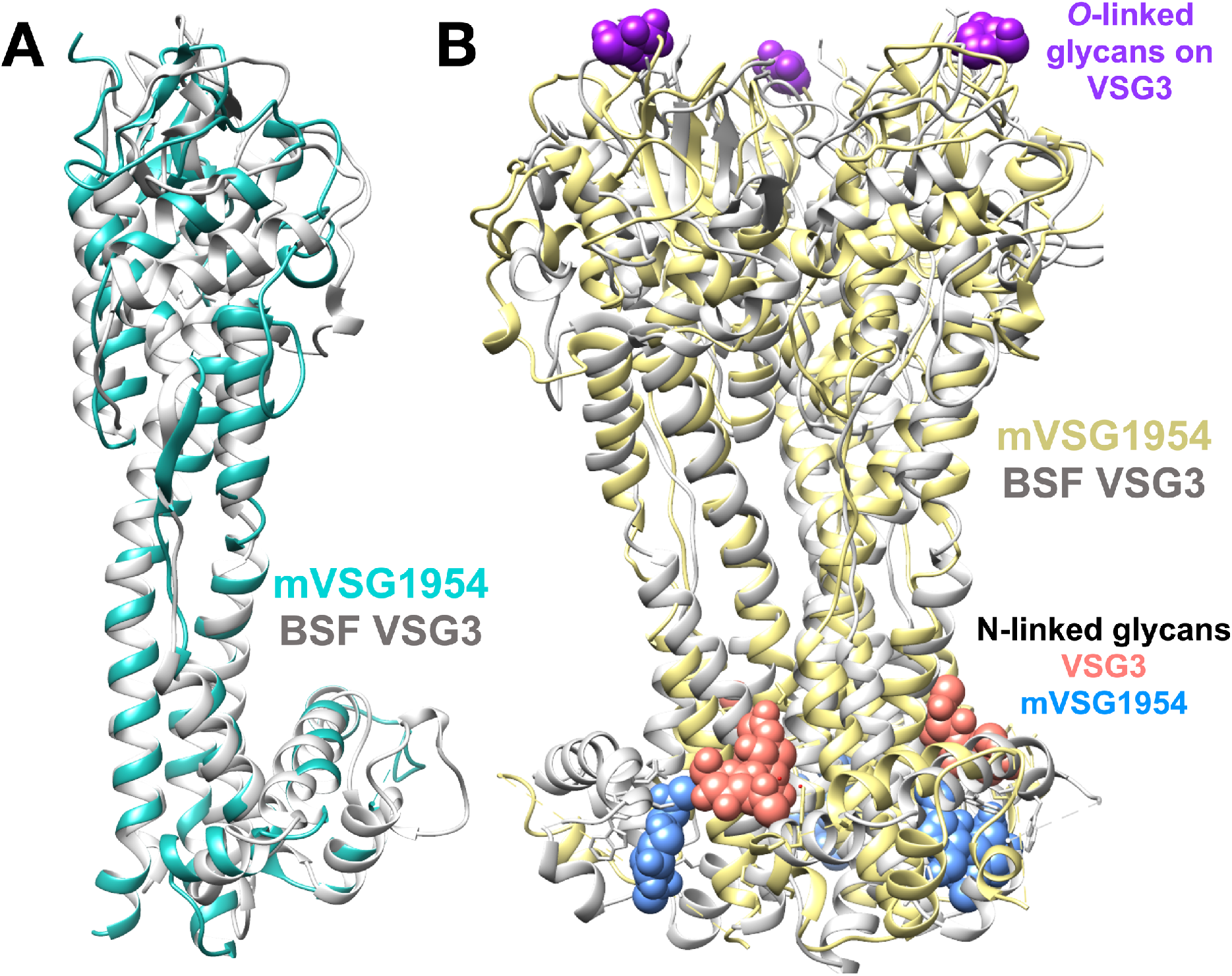
Crystal structure of mVSG1954. (A) Superposition of class B mVSG1954 (aquamarine) and BSF VSG3 (grey) monomers by DeepAlign in RaptorX structure alignment server. (B) Structural superposition of the crystallographic trimers of mVSG1954 (yellow) and VSG3 (grey) performed by SSM Superpose function in COOT (36,63). The *O*-linked sugar on the top lobe of VSG3 is shown in purple as a space-filling atomic representation, whereas the N-linked glycans are shown in salmon (VSG3) and blue (mVSG1954), respectively. Images of protein structures were generated and edited using CHIMERA (61).

As with VSG3, and in contrast to Class A VSGs, mVSG1954 behaves as a monomer in solution (Supplementary Fig 1C) and crystallized as a monomer in the asymmetric unit of the crystal. The ability of mVSG1954 to exist as a monomer is further corroborated by SEC-MALS analysis (size exclusion chromatography coupled to multi-angle light scatting) (20). Also, like VSG3, while a monomer in the asymmetric unit, the mVSG1954 NTD exists as a crystallographic trimer that is highly similar to that of VSG3 (Fig 4B, Supplementary Table S1). Of note is that SEC-MALS data of Class B VSG9 indicates that it forms a trimer in solution (20). Altogether, these data argue that, diverging from the dimers of Class A VSGs, members of Class B may adopt a trimeric oligomeric state, at least in solution and crystals, and possibly on the membrane.

In contrast to the structure of VSG3, mVSG1954 does not show any evidence in the crystals of *O*-linked glycosylation, despite possessing a close consensus sequence to the motif shown previously to often contain this PTM (16). Using intact protein mass spectrometry, the absence of such a sugar in the crystals was confirmed (Supplementary Fig 5). However, because VSG3 sugars have been shown to be heterogeneous (16) and labile (49), it cannot be ruled out that carbohydrate modifications were lost during the process of purification, crystallization, and/or preparation for mass spectrometry.

### Confirming VSG Similarities at the Biological Level

To solve the structures of mVSGs described above, protein crystals were made from mVSGs cloned into BES and purified from the rapid-growing bloodstream form. The structures determined showed a high degree of similarity to their BSF counterparts, but the requirement that the mVSGs be expressed from a BSF expression sites complicates interpretations. This limitation is due to the fact that the metacyclic form of the parasite cannot be effectively cultured as MCFs are cell-cycle arrested. Therefore, the yield of purified mVSGs, especially for the purposes of structural biology and other biochemical examination, is far too low from any such differentiated cells. The need for caution exists because it is possible that cellular machineries associated with the metacyclic stage and the MES could render changes in the mVSGs relative to the BSFs that would be missed by mVSG expression in the BSF stage and expression sites. Such changes could involve PTMs, polypeptide processing, or even conformational differences. For example, in the proteomics data described in Christiano et. al. (29), mVSG653 was reported to be phosphorylated at four amino acids. Phosphorylation has not yet been observed in any available structures of bloodstream VSGs.

To examine this issue, we sought to compare the immunological response to mVSGs as produced in the BSF versus MCF. This process involved (1) producing antisera to mVSGs expressed in the BSFs and then (2) testing metacyclic cultures against these antisera to determine whether they were capable of recognizing mVSGs in the metacyclic context. If the antisera from BSF expressed mVSGs recognize the metacyclic stage, it provides an argument that any changes that might be present between the two forms of the mVSGs produced do not have major immunological consequences. Selecting a member from each of the A and B VSG classes, antisera were raised against both mVSG531 and mVSG1954 expressed in the bloodstream forms in mice (Methods). Metacyclics are cell-cycle arrested and thus cannot be cultured. To obtain metacyclic cells, we differentiated the procyclic form (PCF) of *T. brucei* to the metacyclic form in culture by overexpression of RBP6 protein using tetracycline-inducible system (Methods, Supplementary Figs 6 and 7, (29,50)).

As differentiation proceeds in the fly, the early procyclic surface that is covered in GPEET repeat proteins (a procyclin protein containing Glu-Pro-Glu-Glu-Thr amino acid repeats) is replaced by an EP repeat protein coat (procyclin containing Glu-Pro repeats) in late procyclic cells (51). The Lister 427 PCF cells that we propagate in vitro most likely represent a transition state between these two procyclic cell subtypes, in that they express both GPEET and EP proteins on their coat (Supplementary Fig 6). As these then differentiate, they also lose the EP repeat coat (Supplementary Fig 6). Therefore, we used antibodies against the GPEET and EP to classify the different populations over the course of RBP6 induction. In the starting population, which is the uninduced culture, 91.4% of the cells expressed GPEET on their surface (Supplementary Fig 6A), and 96.6% of the cells express EP on the surface (Supplementary Fig 6B). 98% of the cells are positive when stained with both anti-GPEET and anti-EP, indicated by the shift in fluorescence intensity (Supplementary Fig 6C). We therefore concluded that the majority of the starting population are PCF cells. In the induced culture, 89.9% of the cells had lost their GPEET protein from the surface on day six after induction compared to the starting population (Supplementary Fig 6D), whereas 86.1% of the cells were positive for EP, indicated by the shift in the fluorescence intensity (Supplementary Fig 6E). Only 9.01% of the mixture’s cells did not express either GPEET or EP on the surface (Supplementary Fig 6F), and these likely consist of a combination of epimastigotes (which express BARP), and MCF cells expressing all eight possible Lister 427 mVSGs.

To enrich for differentiated MCFs, cells which still expressed GPEET and EP were separated from the cell mixture by magnetic separation (Methods). The mixture was first stained with both anti-GPEET and anti-EP and counterstained with goat anti-mouse IgG coupled to magnetic beads. The stained PCF cells were subjected to a magnetic field to retain them in the column and separate them from the GPEET and EP negative differentiated cells, which contain an enriched population of the MCF cells of interest (roughly 7 × 10^6^ cells) and epimastigotes (Methods).

The possibility that the metacyclic VSG antisera recognize any marker on the surface of PCFs, which did not cross react with non-cognate BSF cells, also need to be excluded. Therefore, BSF mVSG531 or BSF mVSG1954 were mixed with undifferentiated PCF cells in 1:40 BSF cells to PCF cells ratio. The mixtures were stained with anti-mVSG531 or anti-mVSG1954 and counterstained with anti-mouse IgM conjugated to FITC. The stained cells were subsequently analyzed by flow cytometry. From flow cytometry data, both anti-mVSG531 and anti-mVSG1954 were able to recognize and bind only the small population of BSF cells (presumably expressing mVSG531 or mVSG1954), indicated by 2-3% of cells that has an increased shift in fluorescence intensity (Supplementary Fig 7). This not only demonstrates the lack of cross-reactivity with PCFs, but it also shows that small percentages of cells can be recognized easily by a shift in fluorescence intensity (here, the “spiked” BSF material, comprising only 2-3% of the mix can be identified with ease).

mVSG531 and mVSG1954 primary antisera were used in flow cytometry (in conjunction with a FITC-labeled anti-mouse IgM secondary) to detect mVSG531 and mVSG1954 expressing cells within the purified differentiated MCF population (Methods). Both antisera were first tested to determine whether they could recognize BSF mVSG531 or BSF mVSG1954 (used to infect the mice). For this, BSF mVSG531 or mVSG1954 were incubated with anti-mVSG531 and anti-mVSG1954 and subsequently counterstained using goat anti-mouse IgM conjugated to FITC. The cells were examined using flow cytometry. Both anti-mVSG531 and anti-mVSG1954 recognized the BSF cells expressing their corresponding cognate mVSGs, as indicated by the shift in fluorescence intensity compared to the negative controls, with unstained cells and BSF cells stained with antisera from naïve mice (Fig 5 and Supplementary Fig 8).

**Fig 5:**
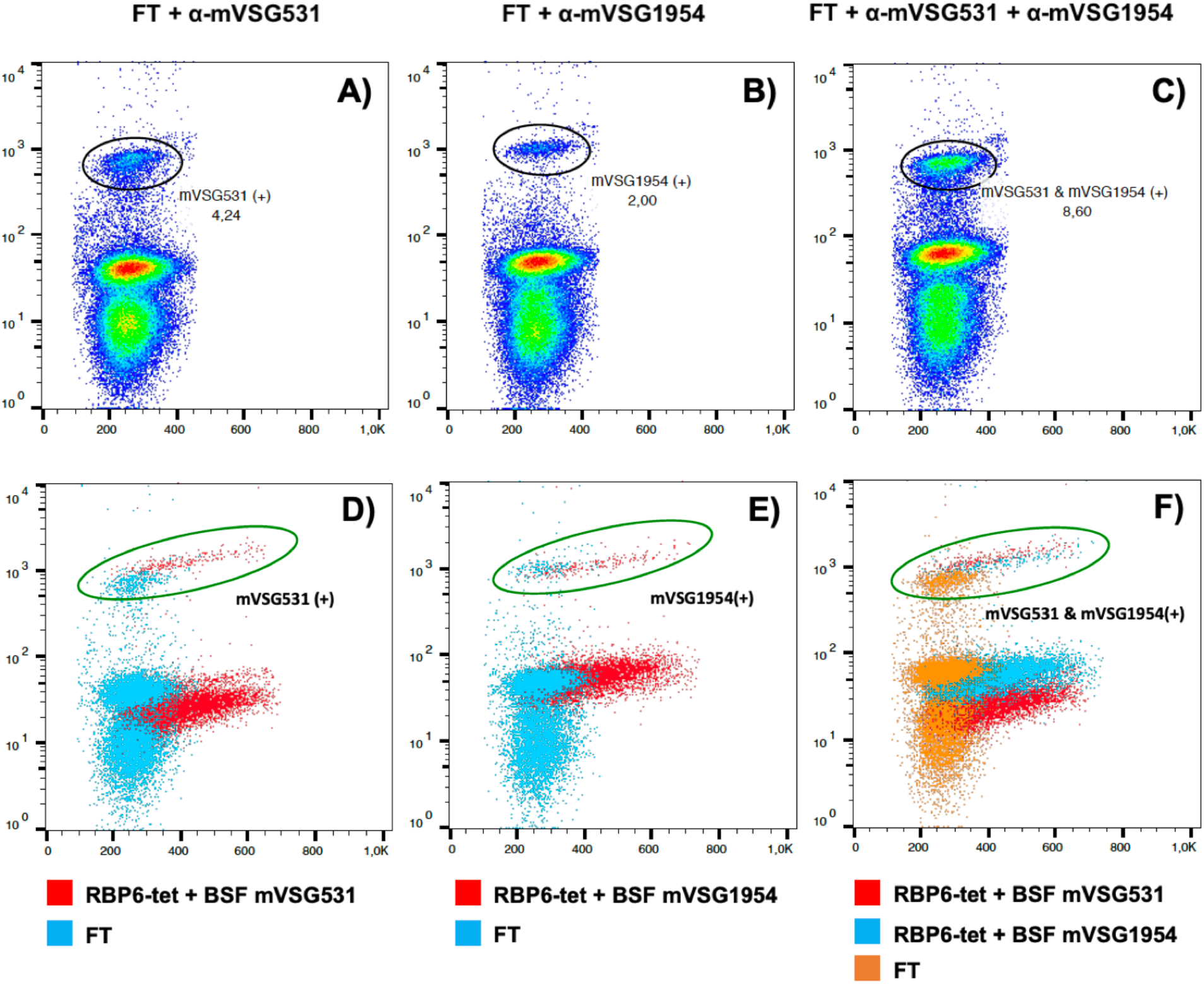
Immune responses to metacyclic and bloodstream expressed VSGs are similar. Antisera to mVSG531 or mVSG1954 elicited in C57BL6/J mice can recognize MCF cells. The undifferentiated PCF cells in the induced culture were separated from the differentiated cells by magnetic labelling. The flow-through fraction that contains the differentiated cells was stained with anti-mVSG531 (A), anti-mVSG1954 (B), or both (C). The cells were further counterstained with goat anti-mouse IgM coupled with FITC. 3.83% and 1.90% cells were positive (A and B, black circle) for either mVSG531 or mVSG1954 on the cells surface, indicated by the shift in the fluorescence intensity. When the cells were stained with both anti-mVSG531 and anti-mVSG1954, 8.18% of the cells were positive (C, black circle). Panels D, E, and F: the flow cytometry dot plots from the differentiated cells overlaid with the dot plot from Supplementary Fig 6 to compare the binding of anti-mVSG531 or anti-mVSG1954 to the cells expressing mVSG531 or mVSG1954 (green circles). “FT” indicates the flow-through fraction that contains differentiated cells.

Additionally, both antisera were tested for cross-reactivity: anti-mVSG531 was tested against BSF mVSG1954 and BSF cells expressing VSG2 (BSF VSG2), while anti-mVSG1954 was tested against BSF mVSG531 and BSF VSG2 as well. No cross-reactivity was detected, and each antibody was only able to recognize the cell type against which each was raised (Supplementary Fig. 8). Therefore, both anti-mVSG531 and anti-mVSG1954 can specifically and exclusively recognize the BSF cells expressing the corresponding mVSGs used in the infection.

Finally, the purified, differentiated MCF cells were stained with anti-mVSG531, anti-mVSG1954, or both (Fig 5). The cells were counter stained with anti-mouse IgM conjugated to FITC and analyzed by flow cytometry. Approximately 3.83% of the cells in the cell mixture were recognized by the anti-mVSG531 antisera, and the level of fluorescence intensity was identical between these MCFs and mVSG531 expressing BSFs (Fig 5, A and D). Similar results were collected for the 1.90% of cells that were recognized by the anti-mVSG1954 antisera (Fig 5, B and E).

Our data suggests that the antisera raised are able to specifically recognize the two mVSGs regardless of whether they were expressed natively in MCFs or heterologously in BSFs. We therefore concluded that lack of an obvious difference in antigenicity implies that, at least with respect to observable immune recognition, these VSGs are identical.

## Discussion

The evolution of the host antibody response and parasite immune evasion mechanisms is a continuous arms race. *T. brucei* overcomes host immunity during the course of infection through a repetitive process of switching VSGs to antigenically distinct versions. The large archive of VSG genes available in *T. brucei’s* genome thereby enables extended survival in the host by continuous antigenic variation (52,53). Aside from amino acid sequence diversity, recent literature has shown that VSGs are also diverse in their tertiary structure organization, oligomerization states, and post-translational modification, factors that can modulate the immune response.

This study focuses on the metacyclic life stage of *T. brucei* in which it inhabits the tsetse fly’s salivary gland and is subsequently transmitted into the mammalian host when the fly takes a blood meal. Like BSF cells, MCF cells also express VSGs on their surface (54). When injected into the mammalian host, the mVSGs expressed on the parasites’ surfaces are retained up to seven days before differentiating into BSF cells that express VSGs from telomeric BESs (55,56). In the Lister 427 *T. brucei* lab model strain used broadly in the study of the African trypanosome, there have been eight VSGs identified that reside in MESs and which are therefore expressed in MCF cells, but structural and functional information on mVSGs has been lacking, particularly whether this unique life cycle stage presented a coat with intrinsic differences from those in the BSF.

This work begins to address these questions at the structural level. The three structures presented here cover a broad range of VSG types, including two divergent Class A molecules and one Class B. Although the metacyclic stage presents unique challenges to the pathogen, these results indicate that mVSGs nonetheless adopt similar three-dimensional properties as their bloodstream counterparts. Furthermore, we show that despite expressing the mVSGs in BES, they appear to be immunologically similar to protein expressed in the metacyclic stage of the parasite.

Altogether, these data suggest that the mVSGs are likely not a specialized subclass of VSG proteins, but instead are derived from the vast genomic archive available through various gene recombination events. This is supported by the fact that the identity of the mVSGs, and even their number, varies dramatically between trypanosome strains. Barring the occurrence of specialized PTMs or other alterations not detected in the BES-produced proteins in this study, the VSG component of the metacyclic coat is not radically different than the BSF.

As MCF cells are the first antigenic surface presented to the mammalian host, and as their mVSG repertoire is limited (eight mVSGs in Lister 427), mVSGs might appear to be an attractive target in the context of prophylaxis treatment (e.g. a vaccine). However, the findings herein that an mVSG itself is “just another” VSG both structurally and antigenically, together with previous studies that have identified mVSG-like proteins expressed within BESs in populations of VSG switchers (57), suggests that the repertoire of mVSGs (or rather, of MES-resident VSGs expressed within MCFs) in nature can be (or can become) just as diverse as that of BES-expressed variants. It is therefore unlikely that mVSGs vaccines can be an effective prophylactic proposition in the context of trypanosomiasis.

## ACKNOWLEDGEMENTS

We acknowledge synchrotron time at the at the Paul Scherrer Institut, Villingen, Switzerland (SLS, beamline PXIII, Vincent Olieric and colleagues) and at the European Synchrotron Radiation Facility (ESRF) in Grenoble (beamline ID29, Gianluca Santoni and colleagues). L.G. was supported by French Government Agence Nationale de la Recherche, ANR-17-CE12-0012 VSGREG. The work was also supported by funds and resources from the German Cancer Research Center (DKFZ) to C.E.S. and F.N.P.

## SUPPLEMENTARY FIGS AND TABLES

Figs 1-8

Table S1

## AUTHOR CONTRIBUTIONS

F.A.B. and M.C. cloned and created trypanosome expressing strains of mVSG397, mVSG531, and mVSG1954. M.v.S. verified mVSG397 clones. M.C. purified and crystallized mVSG531 and mVSG1954, solved and refined their crystal structures (collecting data on mVSG1954 with J.Z.), generated their antisera, conducted trypanosome differentiation and FACs experiments with F.N.P., co-wrote the manuscript, and generated figures. J.Z. and K.F. expressed, purified, and crystallized mVSG397 while J.Z. and S.D. collected crystallographic data and solved the structure of mVSG397. S.D. conducted suramin survival experiments and S.D. and M.v.S. analyzed and graphed this data. E.T. optimized the trypanosome differentiation protocol and provided reagents and L.G. edited the manuscript. N.L. and T.R. performed and analyzed mass spectrometry experiments. C.E.S. conceived of the projects, aided in structural determination, co-wrote the manuscript, and generated figures. All authors commented and contributed to the writing of the manuscript.

## Data Availability

Coordinates and structural factors have been uploaded to the RCSB PDB (www.rcsb.org): mVSG397 (PDB ID 8B3E), mVSG531 (PDB ID 8B3B), and mVSG1954 (PDB ID 8B3W). Other data supporting the findings of this study are available from the authors upon request.

## REFERENCES

1. Keating J, Yukich JO, Sutherland CS, Woods G, Tediosi F. Human African trypanosomiasis prevention, treatment and control costs: A systematic review. Acta Trop. 2015 Oct 1;150:4–13.

2. Ponte-Sucre A. An Overview of Trypanosoma brucei Infections: An Intense Host–Parasite Interaction. Front Microbiol [Internet]. 2016 Dec 26 [cited 2020 Feb 18];7. Available from: http://journal.frontiersin.org/article/10.3389/fmicb.2016.02126/full

3. Matthews KR, McCulloch R, Morrison LJ. The within-host dynamics of African trypanosome infections. Philos Trans R Soc Lond B Biol Sci. 2015 Aug 19;370(1675).

4. Bangs JD. Evolution of Antigenic Variation in African Trypanosomes: Variant Surface Glycoprotein Expression, Structure, and Function. BioEssays. 2018;40(12):1800181.

5. Manna PT, Boehm C, Leung KF, Natesan SK, Field MC. Life and times: synthesis, trafficking, and evolution of VSG. Trends Parasitol. 2014 May;30(5):251–8.

6. Borst P, Rudenko G, Blundell PA, van Leeuwen F, Cross MA, McCulloch R, et al. Mechanisms of antigenic variation in African trypanosomes. Behring Inst Mitt. 1997 Mar;(99):1–15.

7. Horn D. Antigenic variation in African trypanosomes. Mol Biochem Parasitol. 2014 Jul;195(2):123–9.

8. Mugnier MR, Stebbins CE, Papavasiliou FN. Masters of Disguise: Antigenic Variation and the VSG Coat in Trypanosoma brucei. PLOS Pathog. 2016 Sep 1;12(9):e1005784.

9. Aresta-Branco F, Erben E, Papavasiliou FN, Stebbins CE. Mechanistic Similarities between Antigenic Variation and Antibody Diversification during Trypanosoma brucei Infection. Trends Parasitol. 2019;35(4):302–15.

10. Cross GA, Kim HS, Wickstead B. Capturing the variant surface glycoprotein repertoire (the VSGnome) of Trypanosoma brucei Lister 427. Mol Biochem Parasitol. 2014 Jun;195(1):59–73.

11. Marcello L, Barry JD. Analysis of the VSG gene silent archive in Trypanosoma brucei reveals that mosaic gene expression is prominent in antigenic variation and is favored by archive substructure. Genome Res. 2007 Sep;17(9):1344–52.

12. Weirather JL, Wilson ME, Donelson JE. Mapping of VSG similarities in Trypanosoma brucei. Mol Biochem Parasitol. 2012 Feb;181(2):141–52.

13. Bartossek T, Jones NG, Schafer C, Cvitkovic M, Glogger M, Mott HR, et al. Structural basis for the shielding function of the dynamic trypanosome variant surface glycoprotein coat. Nat Microbiol. 2017 Nov;2(11):1523–32.

14. Carrington M, Miller N, Blum M, Roditi I, Wiley D, Turner M. Variant specific glycoprotein of Trypanosoma brucei consists of two domains each having an independently conserved pattern of cysteine residues. J Mol Biol. 1991 Oct 5;221(3):823–35.

15. Freymann D, Down J, Carrington M, Roditi I, Turner M, Wiley D. 2.9 A resolution structure of the N-terminal domain of a variant surface glycoprotein from Trypanosoma brucei. J Mol Biol. 1990 Nov 5;216(1):141–60.

16. Pinger J, Nešić D, Ali L, Aresta-Branco F, Lilic M, Chowdhury S, et al. African trypanosomes evade immune clearance by O -glycosylation of the VSG surface coat. Nat Microbiol. 2018 Aug;3(8):932–8.

17. Zeelen J, van Straaten M, Verdi J, Hempelmann A, Hashemi H, Perez K, et al. Structure of trypanosome coat protein VSGsur and function in suramin resistance. Nat Microbiol. 2021 Mar;6(3):392–400.

18. Chattopadhyay A, Jones NG, Nietlispach D, Nielsen PR, Voorheis HP, Mott HR, et al. Structure of the C-terminal domain from Trypanosoma brucei variant surface glycoprotein MITat1.2. J Biol Chem. 2005 Feb 25;280(8):7228–35.

19. Jones NG, Nietlispach D, Sharma R, Burke DF, Eyres I, Mues M, et al. Structure of a glycosylphosphatidylinositol-anchored domain from a trypanosome variant surface glycoprotein. J Biol Chem. 2008 Feb 8;283(6):3584–93.

20. Umaer K, Aresta-Branco F, Chandra M, van Straaten M, Zeelen J, Lapouge K, et al. Dynamic, variable oligomerization and the trafficking of variant surface glycoproteins of Trypanosoma brucei. Traffic Cph Den. 2021 Aug;22(8):274–83.

21. Alarcon CM, Son HJ, Hall T, Donelson JE. A monocistronic transcript for a trypanosome variant surface glycoprotein. Mol Cell Biol. 1994 Aug;14(8):5579–91.

22. Graham SV, Barry JD. Transcriptional regulation of metacyclic variant surface glycoprotein gene expression during the life cycle of Trypanosoma brucei. Mol Cell Biol. 1995 Nov;15(11):5945–56.

23. Hutchinson S, Foulon S, Crouzols A, Menafra R, Rotureau B, Griffiths AD, et al. The establishment of variant surface glycoprotein monoallelic expression revealed by single-cell RNA-seq of Trypanosoma brucei in the tsetse fly salivary glands. Deitsch KW, editor. PLOS Pathog. 2021 Sep 20;17(9):e1009904.

24. Müller LSM, Cosentino RO, Förstner KU, Guizetti J, Wedel C, Kaplan N, et al. Genome organization and DNA accessibility control antigenic variation in trypanosomes. Nature. 2018 Nov;563(7729):121–5.

25. Brun R, Blum J, Chappuis F, Burri C. Human African trypanosomiasis. Lancet Lond Engl. 2010 Jan 9;375(9709):148–59.

26. Büscher P, Cecchi G, Jamonneau V, Priotto G. Human African trypanosomiasis. The Lancet. 2017 Nov;390(10110):2397–409.

27. Matthews KR. The developmental cell biology of Trypanosoma brucei. J Cell Sci. 2005 Jan 15;118(Pt 2):283–90.

28. Romero-Meza G, Mugnier MR. Trypanosoma brucei. Trends Parasitol. 2020 Jun;36(6):571–2.

29. Christiano R, Kolev NG, Shi H, Ullu E, Walther TC, Tschudi C. The proteome and transcriptome of the infectious metacyclic form of Trypanosoma brucei define quiescent cells primed for mammalian invasion. Mol Microbiol. 2017 Oct;106(1):74–92.

30. Alsford S, Kawahara T, Glover L, Horn D. Tagging a T. brucei RRNA locus improves stable transfection efficiency and circumvents inducible expression position effects. Mol Biochem Parasitol. 2005 Dec;144(2):142–8.

31. Cross GA. Release and purification of Trypanosoma brucei variant surface glycoprotein. J Cell Biochem. 1984;24(1):79–90.

32. Ness SR, de Graaff RAG, Abrahams JP, Pannu NS. CRANK: new methods for automated macromolecular crystal structure solution. Struct Lond Engl 1993. 2004 Oct;12(10):1753–61.

33. Cowtan K. Completion of autobuilt protein models using a database of protein fragments. Acta Crystallogr D Biol Crystallogr. 2012 Apr 1;68(4):328–35.

34. Adams PD, Afonine PV, Bunkoczi G, Chen VB, Davis IW, Echols N, et al. PHENIX: a comprehensive Python-based system for macromolecular structure solution. Acta Crystallogr Biol Crystallogr. 2010 Feb;66(Pt 2):213–21.

35. Liebschner D, Afonine PV, Baker ML, Bunkóczi G, Chen VB, Croll TI, et al. Macromolecular structure determination using X-rays, neutrons and electrons: recent developments in Phenix. Acta Crystallogr Sect Struct Biol. 2019 Oct 1;75(10):861–77.

36. Emsley P, Cowtan K. Coot: model-building tools for molecular graphics. Acta Crystallogr D Biol Crystallogr. 2004 Dec 1;60(12):2126–32.

37. Joosten RP, Long F, Murshudov GN, Perrakis A. The PDB_REDO server for macromolecular structure model optimization. IUCrJ. 2014 Jul 1;1(Pt 4):213–20.

38. McCoy AJ, Grosse-Kunstleve RW, Adams PD, Winn MD, Storoni LC, Read RJ. Phaser crystallographic software. J Appl Crystallogr. 2007 Aug 1;40(4):658–74.

39. Terwilliger TC, Grosse-Kunstleve RW, Afonine PV, Moriarty NW, Zwart PH, Hung LW, et al. Iterative model building, structure refinement and density modification with the PHENIX AutoBuild wizard. Acta Crystallogr D Biol Crystallogr. 2008 Jan 1;64(1):61–9.

40. Sheldrick GM. A short history of SHELX. Acta Crystallogr A. 2008 Jan;64(Pt 1):112–22.

41. Potterton L, Agirre J, Ballard C, Cowtan K, Dodson E, Evans PR, et al. CCP4i2: the new graphical user interface to the CCP4 program suite. Acta Crystallogr Sect Struct Biol. 2018 Feb 1;74(2):68–84.

42. Langer G, Cohen SX, Lamzin VS, Perrakis A. Automated macromolecular model building for X-ray crystallography using ARP/wARP version 7. Nat Protoc. 2008;3(7):1171–9.

43. Potterton L, Agirre J, Ballard C, Cowtan K, Dodson E, Evans PR, et al. CCP 4 i 2: the new graphical user interface to the CCP 4 program suite. Acta Crystallogr Sect Struct Biol. 2018 Feb 1;74(2):68–84.

44. Winter G. xia2 : an expert system for macromolecular crystallography data reduction. J Appl Crystallogr. 2010 Feb 1;43(1):186–90.

45. Winter G, Beilsten-Edmands J, Devenish N, Gerstel M, Gildea RJ, McDonagh D, et al. DIALS as a toolkit. Protein Sci Publ Protein Soc. 2022 Jan;31(1):232–50.

46. Tihon E, Rubio-Peña K, Dujeancourt-Henry A, Crouzols A, Rotureau B, Glover L. VEX1 Influences mVSG Expression During the Transition to Mammalian Infectivity in Trypanosoma brucei. Front Cell Dev Biol. 2022;10:851475.

47. Wang S, Ma J, Peng J, Xu J. Protein structure alignment beyond spatial proximity. Sci Rep. 2013;3:1448.

48. Wang S, Peng J, Xu J. Alignment of distantly related protein structures: algorithm, bound and implications to homology modeling. Bioinforma Oxf Engl. 2011 Sep 15;27(18):2537–45.

49. Gkeka A, Aresta-Branco F, Triller G, Vlachou EP, Lilic M, Olinares PDB, et al. Immunodominant surface epitopes power immune evasion in the African trypanosome [Internet]. 2021 Jul [cited 2021 Sep 21] p. 2021.07.20.453071. Available from: https://www.biorxiv.org/content/10.1101/2021.07.20.453071v1

50. Kolev NG, Ramey-Butler K, Cross GAM, Ullu E, Tschudi C. Developmental progression to infectivity in Trypanosoma brucei triggered by an RNA-binding protein. Science. 2012 Dec 7;338(6112):1352–3.

51. Acosta-Serrano A, Vassella E, Liniger M, Renggli CK, Brun R, Roditi I, et al. The surface coat of procyclic Trypanosoma brucei: Programmed expression and proteolytic cleavage of procyclin in the tsetse fly. Proc Natl Acad Sci. 2001 Feb 13;98(4):1513–8.

52. Hovel-Miner G, Mugnier M, Papavasiliou FN, Pinger J, Schulz D. A Host-Pathogen Interaction Reduced to First Principles: Antigenic Variation in T. brucei. Results Probl Cell Differ. 2015;57:23–46.

53. Mugnier MR, Stebbins CE, Papavasiliou FN. Masters of Disguise: Antigenic Variation and the VSG Coat in Trypanosoma brucei. PLoS Pathog. 2016 Sep;12(9):e1005784.

54. Tetley L, Turner CM, Barry JD, Crowe JS, Vickerman K. Onset of expression of the variant surface glycoproteins of Trypanosoma brucei in the tsetse fly studied using immunoelectron microscopy. J Cell Sci. 1987 Mar;87 (Pt 2):363–72.

55. Graham SV, Matthews KR, Shiels PG, Barry JD. Distinct, developmental stage-specific activation mechanisms of trypanosome VSG genes. Parasitology. 1990 Dec;101(3):361–7.

56. Helm JR, Hertz-Fowler C, Aslett M, Berriman M, Sanders M, Quail MA, et al. Analysis of expressed sequence tags from the four main developmental stages of Trypanosoma congolense. Mol Biochem Parasitol. 2009 Nov;168(1):34–42.

57. Mugnier MR, Cross GA, Papavasiliou FN. The in vivo dynamics of antigenic variation in Trypanosoma brucei. Science. 2015 Mar 27;347(6229):1470–3.

58. McNicholas S, Potterton E, Wilson KS, Noble MEM. Presenting your structures: the CCP4mg molecular-graphics software. Acta Crystallogr D Biol Crystallogr. 2011 Apr 1;67(Pt 4):386–94.

59. Laskowski RA, Jabłońska J, Pravda L, Vařeková RS, Thornton JM. PDBsum: Structural summaries of PDB entries. Protein Sci. 2018 Jan;27(1):129–34.

60. Schrodinger. The PyMOL Molecular Graphics System, Version 1.8. 2015.

61. Pettersen EF, Goddard TD, Huang CC, Couch GS, Greenblatt DM, Meng EC, et al. UCSF Chimera--a visualization system for exploratory research and analysis. J Comput Chem. 2004 Oct;25(13):1605–12.

62. Eisenberg D, Schwarz E, Komaromy M, Wall R. Analysis of membrane and surface protein sequences with the hydrophobic moment plot. J Mol Biol. 1984 Oct 15;179(1):125–42.

63. Emsley P, Lohkamp B, Scott WG, Cowtan K. Features and development of Coot. Acta Crystallogr D Biol Crystallogr. 2010 Apr;66(Pt 4):486–501.

