## Supplementary figures, methods, and tables for "Structural and Immunological Similarities Between the Metacyclic and Bloodstream Form Variant Surface Glycoproteins of the African Trypanosome"

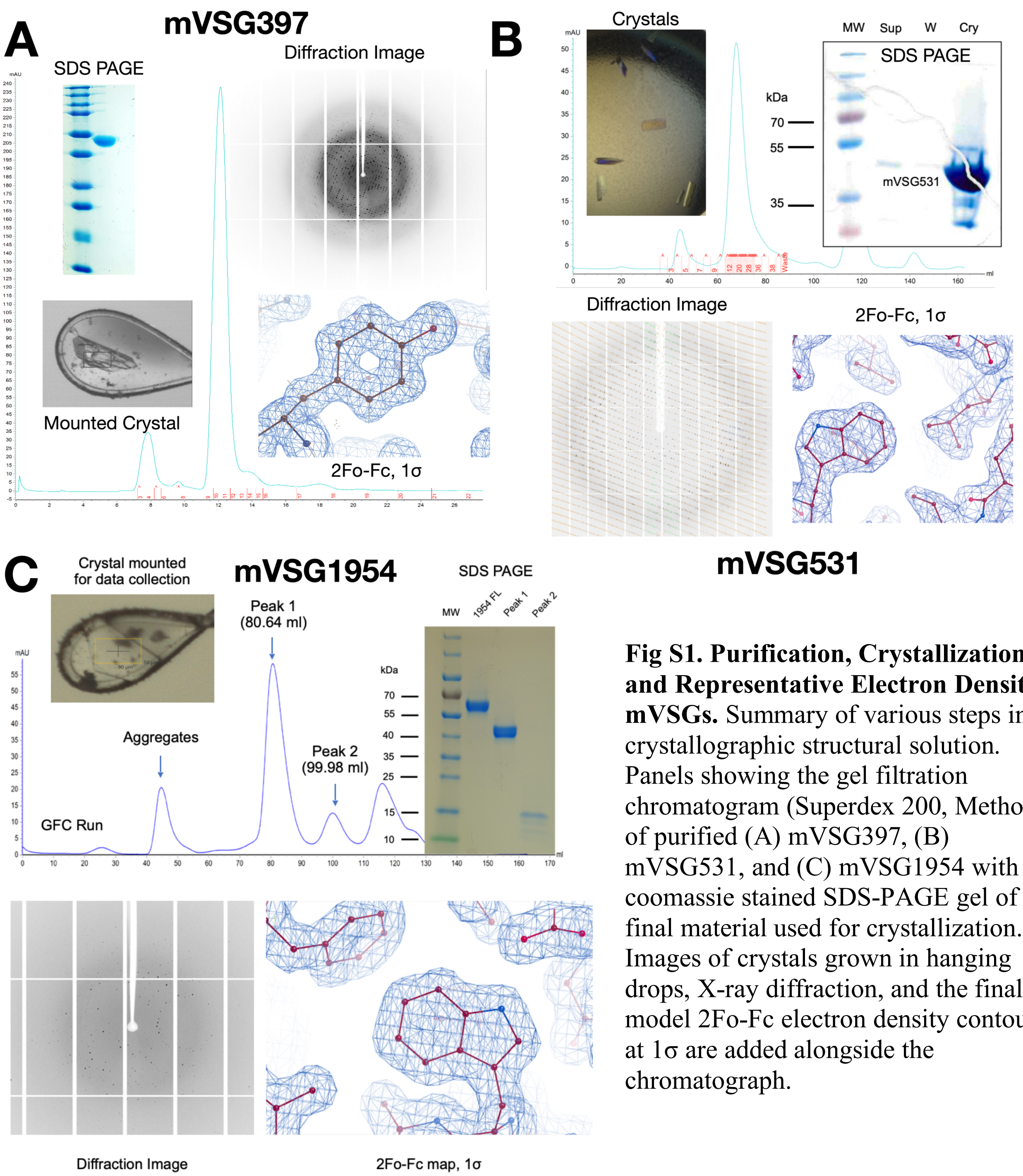


**Fig S1. Purification, Crystallization, and Representative Electron Density of mVSGs.** Summary of various steps in the crystallographic structural solution. Panels showing the gel filtration chromatogram (Superdex 200, Methods) of purified (A) mVSG397, (B) mVSG531, and (C) mVSG1954 with a coomassie stained SDS-PAGE gel of the final material used for crystallization. Images of crystals grown in hanging drops, X-ray diffraction, and the final model 2Fo-Fc electron density contoured at 1σ are added alongside the chromatograph.

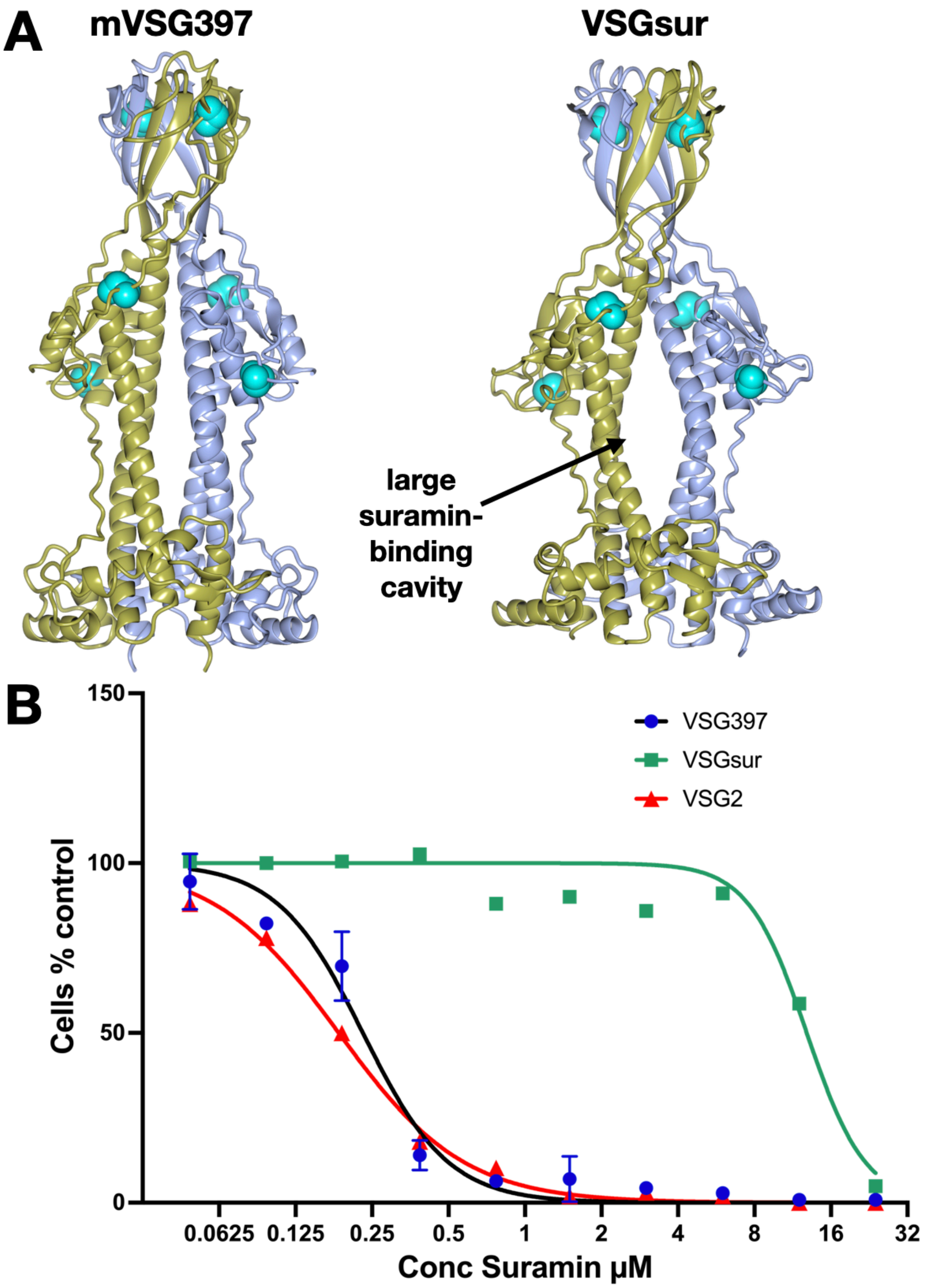


**Fig S2. Structural and Functional Comparisons of mVSG397 with VSGsur**

(A) Ribbon diagrams showing that the pocket between monomers is much smaller in mVSG397 as compared to VSGsur and (B) Suramin resistance assays showing that mVSG397 does not confer resistance to suramin. Differences between VSG397 and VSG2 are not statistically significant (two-tailed P=0.3029) whereas the difference between VSG397 and VSGsur are statistically significant (two-tailed P<0.0001).

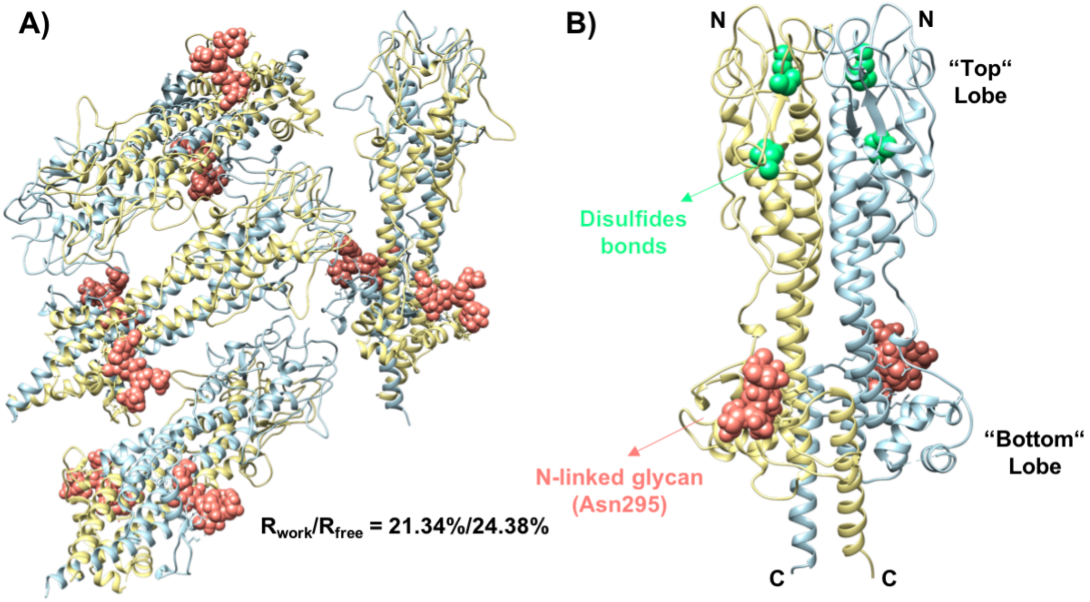


**Fig S3. Atomic resolution structure of mVSG531.** Molecular replacement was performed by PHASER-MR on the native crystal dataset using BSF VSG1 as search model to determine the structure of mVSG531 (Methods). (A) Eight molecules of mVSG531 was found in the crystal asymmetric unit. (B) N-linked glycan is attached at Asn295 at the dimer's bottom lobe (salmon pink spheres). Two disulfide bonds are observed at the top lobe of each molecule (green spheres). Images of protein structures were generated and edited using CHIMERA.

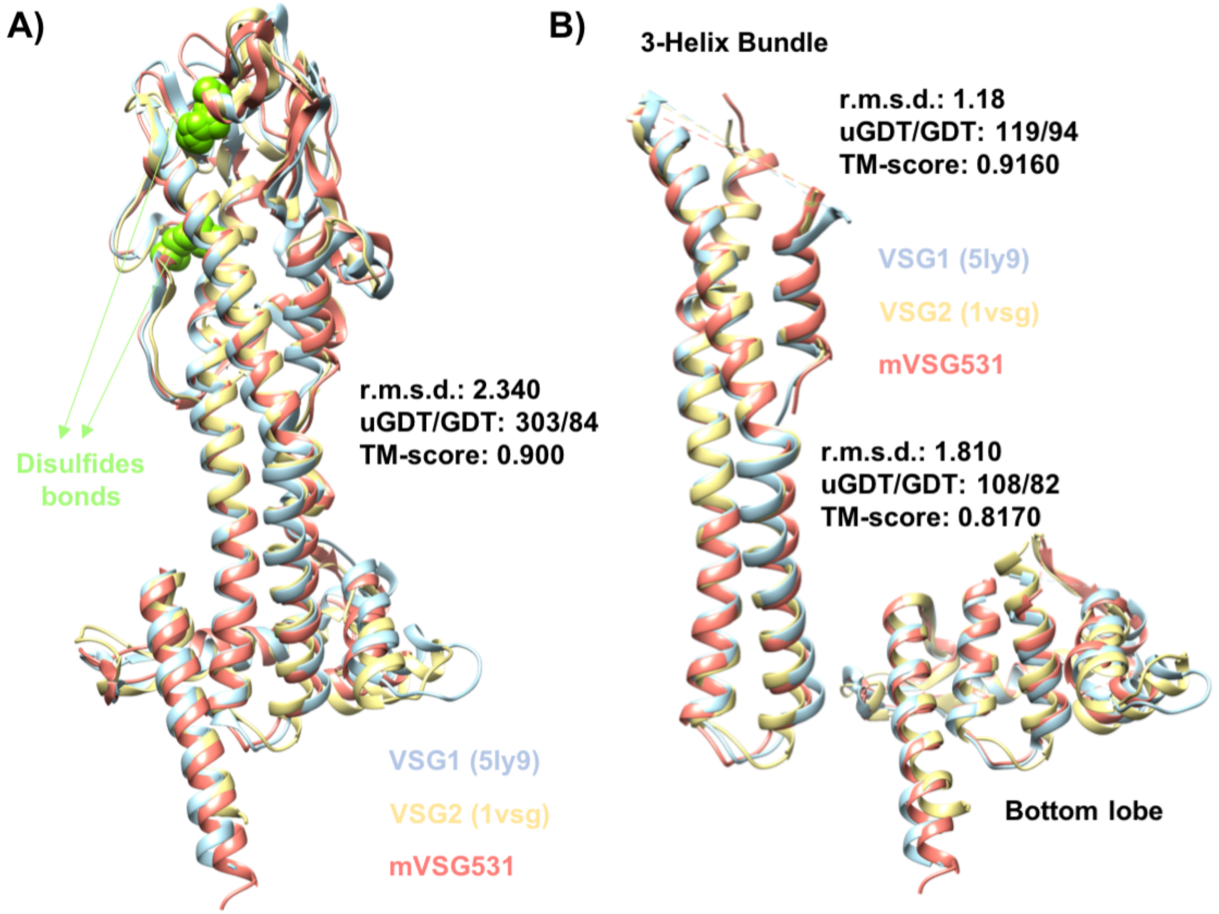


**Fig S4. Structure alignment between mVSG531, VSG2, and VSG1 reveals structure similarity between mVSG531 and bloodstream VSGs in Class A.** (A) Overall structural alignment of mVSG531, VSG2 (PDB code: 1VSG), and VSG1 (PDB code: 5LY9) monomers (yellow: VSG2, blue: VSG1, salmon pink: mVSG531, green: disulfide bonds) (B) Three-helix bundle core (left) and the bottom lobe (right) structural alignment. Structural alignment was performed by DeepAlign in RaptorX structure alignment server. (r.m.s.d.: root mean square deviation uGDT/GDT: (unnormalized) global distance test, TM-score: template modeling score). Images of protein structures were generated using CHIMERA.

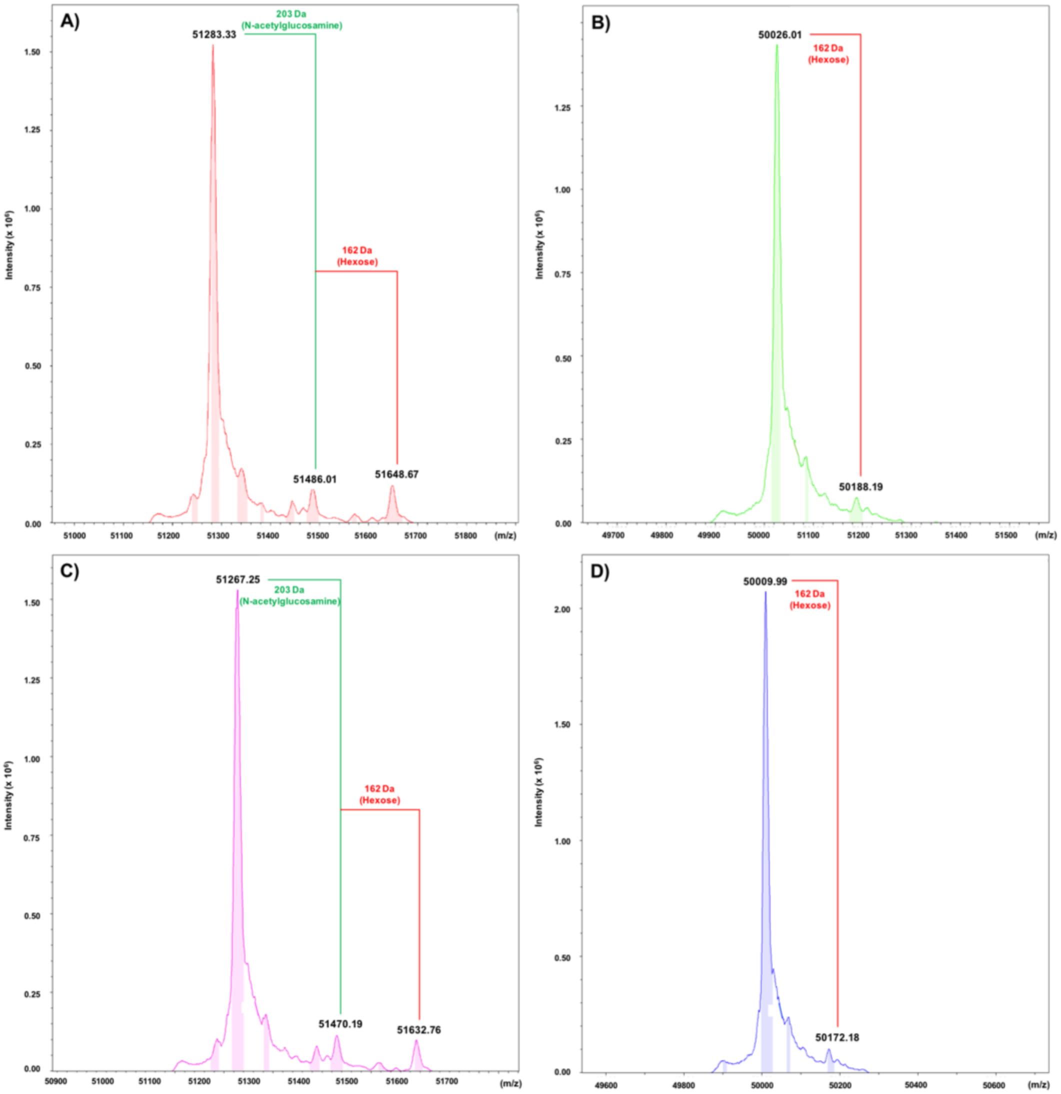


**Fig S5. mVSG1954 shows no O-glycosylation by mass spectrometry.** Intact mass spectrometry was performed on full-length mVSG1954 wildtype and S321A (A and C). Both protein constructs are also treated with PNGaseF to cleave the N-linked glycans (B and D). Mass differences of 203 Da and 162 Da was observed in the full-length mVSG1954 wildtype (A), which correspond to N-acetylglucosamine and hexose molecules, respectively. When the sample was treated with PNGaseF (B), only the 162 Da mass difference was observed, indicating the loss of N-linked glycan and the possible presence of *O*-linked glycan. This was examined by mutating the putative O-glycosylation site at S321 to alanine, which cannot be *O*-glycosylated. A 16 Da mass difference was observed between the full-length mVSG1954 wildtype (A) and S321A (C), indicating the serine to alanine mutation. However, when treated with PNGaseF (D), the 162 Da mass difference still presents, indicating that the protein is not *O*-glycosylated on that residue. Since the material is FL VSG1954, it is possible the loss of a mass equal to a hexose could be from the CTD GPI anchor.

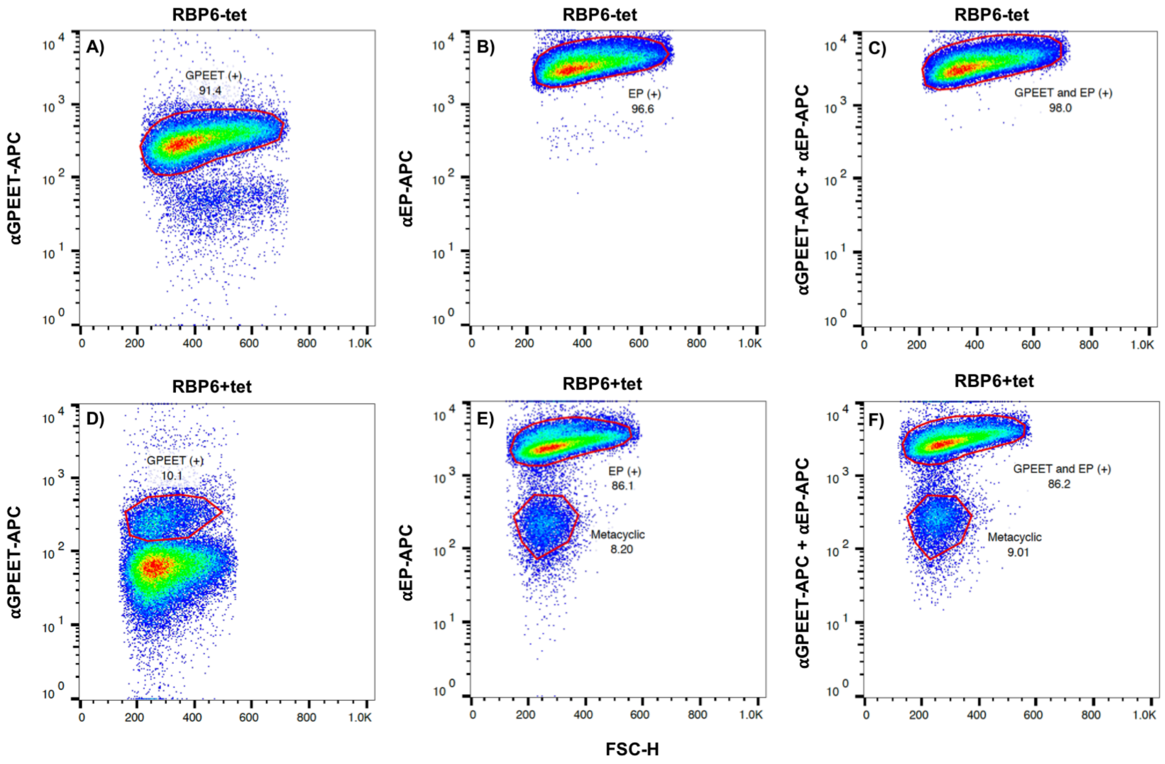


**Fig S6. Overexpression of RBP6 in PCF cells initiate differentiation of PCF cells into later life stages of *T. brucei*.** Wildtype 29-13 PCF cells tetracycline-inducible RBP6 overexpression plasmid was cultured in SDM-80 media supplemented with 7.5 µg/ml Hemin, 50 mM N-Acetylglucosamine, and 10% v/v heat-inactivated FBS at 27oC and 5% $CO_2$ (Methods). The induction was started at 2 million cells/ml cell density with 10 µg/ml tetracycline for 6 days. On day 6, both uninduced and induced culture were analyzed by flow cytometry. Both uninduced and induced cultures were stained with a-GPEET (A, D), a-EP (B, E), or both a-GPEET and a-EP (C, F). Both a-GPEET and a-EP were coupled to fluorescence molecule APC. 91.4% of the cells in starting population (uninduced culture) is positive for GPEET on their cell surface (A), and 96.6% of the cells is positive for EP (B), indicated by the shift in the fluorescence intensity. When stained with both a-GPEET and a-EP, 98% of the starting population has shifted in fluorescence intensity (C). In the induced culture, only 10.1% from the cells is positive for GPEET and 86.1% of the cells is positive for EP. When the induced culture is stained with both a-GPEET and a-EP 86.2% is positive for both GPEET and EP. 9.01% of the cells in the induced culture is negative for either GPEET and EP, indicating that this cell population is no longer PCF cells.

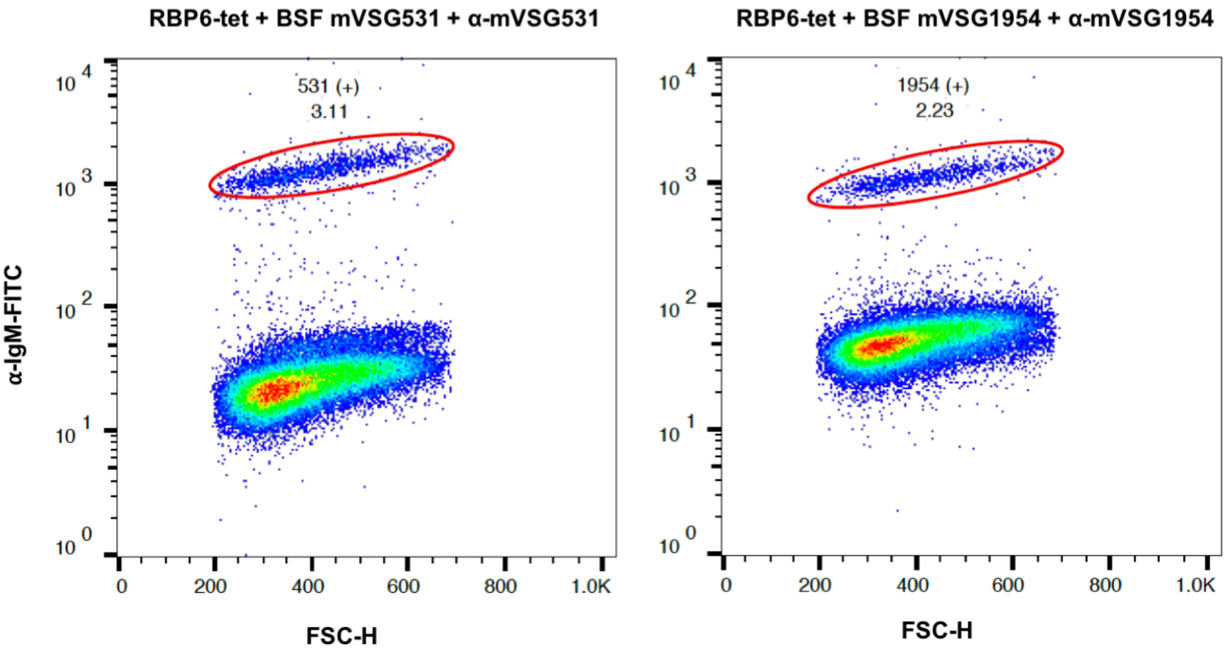


**Fig S7. Antisera a-mVSG531 and a-mVSG1954 elicited in C57BL6/J mice is able to recognize small positive cell population from a mixture with undifferentiated PCF cells.** 5% of BSF cells expressing mVSG531 (left) or mVSG1954 (right) was mixed with 95% undifferentiated PCF cells. The mixtures were stained with either a-mVSG531 or a- mVSG1954 and counterstained with Goat anti-mouse IgM coupled to FITC. 3.11% of the cells in the mixture of undifferentiated PCF cells and BSF cells expressing mVSG531 has shifted in fluorescence intensity (left). 2.33% of the cells in the mixture of undifferentiated PCF cells and BSF cells expressing mVSG1954 has shifted in fluorescence intensity (right).

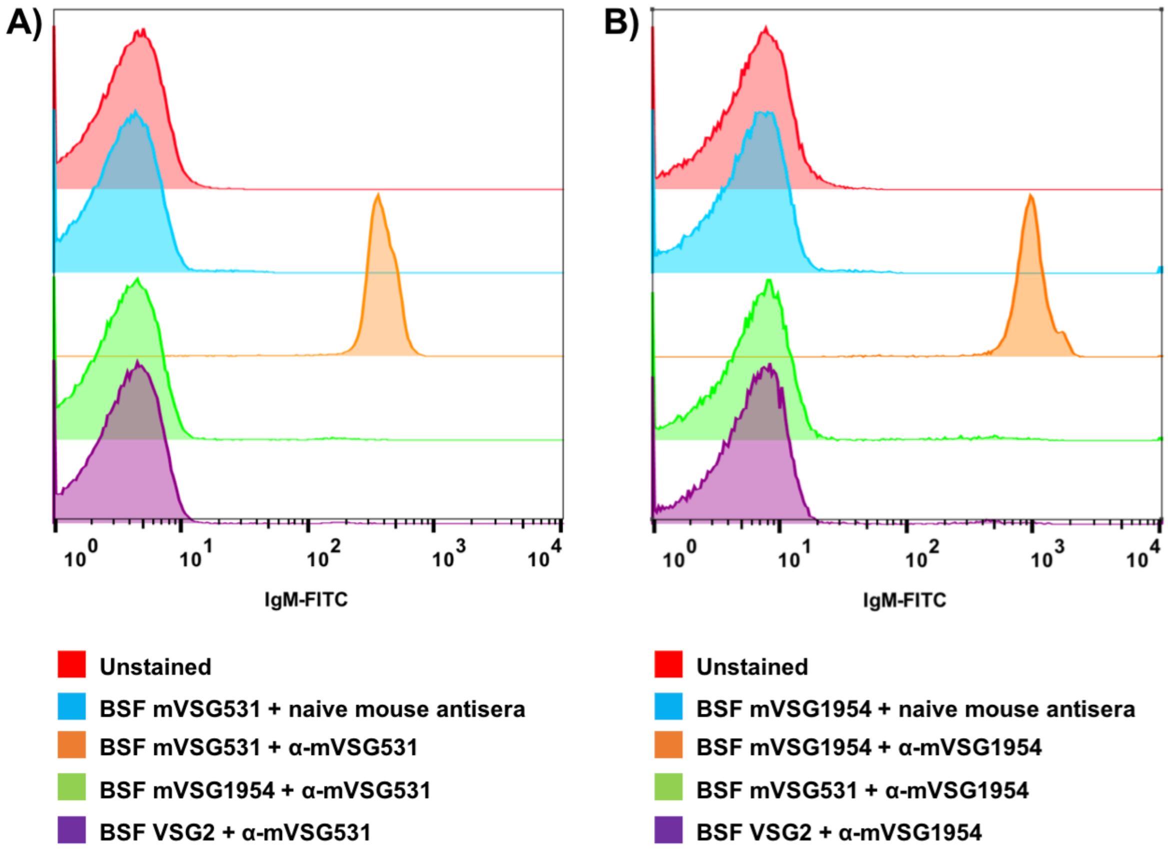


**Fig S8. Antisera raised against mVSG531 or mVSG1954 in C57BL6/J mice specifically recognize the BSF cells expressing respective mVSGs.** (A) anti-mVSG531 was tested to stain BSF mVSG531 (orange), BSF mVSG1954 (green) and BSF VSG2 (purple). Cells were counterstained with secondary goat anti-mouse IgM coupled to FITC. Unstained cells and BSF cells expressing mVSG531 stained with antisera from naïve mouse were used as negative controls. (B) anti-mVSG1954 was tested to stain BSF mVSG1954 (orange), BSF mVSG531 (green) and BSF VSG2 (purple). Cells were counterstained with secondary goat anti-mouse IgM coupled to FITC. Unstained cells and BSF cells expressing mVSG1954 stained with antisera from naïve mouse were used as negative controls. Shift in fluorescence intensity was observed in the sample in which the antisera were used to stain the BSF cells expressing the respective mVSG (orange), but not in the sample in which the antisera were used to stain BSF cells expressing the opposite mVSG (green) or BSF VSG2 (purple).

**Table S1: Crystallographic Statistics**

| Parameter | VSG531 | VSG1954 | VSG397 |
|---|---|---|---|
| Wavelength | 1.00 Å | 0.9198 Å | 1.00 Å |
| Resolution range | 59.09-1.95 (2.021.95) | 60.29-1.68 (1.74-1.68) | 54.62-1.26 (1.305-1.26) |
| Space group | P 1 21 1 | P 3 2 1 | C 2 2 21 |
| Unit cell | 50.107 170.88 163.65 90 90.705 90 | 69.614 69.614 115.36 90 90 120 | 55.56 218.48 66.44 90 90 90 |
| Total reflections | 592350 (57625) | 723113 (71206) | 536522 (24839) |
| Unique reflections | 195472 (18665) | 37543 (3705) | 96793 (8421) |
| Multiplicity | 3.0 (3.1) | 19.3 (19.2) | 5.5 (2.9) |
| Completeness (%) | 97.68 (93.16) | 99.79 (97.89) | 96.37 (79.98) |
| Mean I/sigma(I) | 4.24 (1.13) | 7.45 (0.48) | 6.56 (0.42) |
| Wilson B-factor | 34.64 | 19.13 | 15.19 |
| R-merge | 0.1157 (0.7639) | 0.1453 (1.283) | 0.0879 (1.321) |
| R-meas | 0.1412 (0.9266) | 0.1492 (1.318) | 0.0972 (1.575) |
| R-pim | 0.07975 (0.5179) | 0.03383 (0.2986) | 0.04045( 0.8407) |
| CC1/2 | 0.977 (0.629) | 0.999 (0.772) | 0.99 (0.219) |
| CC* | 0.994 (0.879) | 1 (0.934) | 0.998 (0.599) |
| Reflections (refinement) | 195256 (18624) | 37464 (3627) | 95996 (7883) |
| Reflections (R-free) | 9605 (847) | 1877 (201) | 4807 (404) |
| R-work | 0.2134 (0.3583) | 0.1757 (0.3505) | 0.1834 (0.3799) |
| R-free | 0.2438 (0.3808) | 0.2137 (0.3762) | 0.2160 (0.4115) |
| CC(work) | 0.916 (0.772) | 0.968 (0.829) | 0.960 (0.596) |
| CC(free) | 0.904 (0.745) | 0.971 (0.783) | 0.973 (0.596) |
| Non-hydrogen atoms | 22693 | 2798 | 3366 |
| macromolecules | 20778 | 2514 | 2882 |
| ligands | 618 | 62 | 53 |
| solvent | 1339 | 250 | 456 |
| Protein residues | 2866 | 353 | 389 |
| RMS(bonds) | 0.006 | 0.011 | 0.009 |
| RMS(angles) | 1.16 | 1.34 | 1.21 |
| Ramachandran favored (%) | 96.78 | 98.84 | 98.44 |
| Ramachandran allowed (%) | 3.22 | 1.16 | 1.30 |
| Ramachandran outliers (%) | 0.00 | 0.00 | 0.26 |
| Rotamer outliers (%) | 0.25 | 0.41 | 0.68 |
| Clashscore | 1.81 | 3.19 | 1.55 |
| Average B-factor | 46.61 | 29.42 | 25.88 |
| macromolecules | 47.27 | 28.74 | 24.30 |
| ligands | 30.97 | 61.13 | 67.38 |
| solvent | 43.09 | 31.97 | 33.92 |
| Number of TLS groups | 45 | 5 | 5 |

Statistics for the highest-resolution shell are shown in parentheses.
